# Polychaetoid/ZO-1 strengthens cell junctions under tension while localizing differently than core adherens junction proteins

**DOI:** 10.1101/2023.03.01.530634

**Authors:** Anja Schmidt, Tara Finegan, Matthias Häring, Deqing Kong, Alexander G Fletcher, Zuhayr Alam, Jörg Grosshans, Fred Wolf, Mark Peifer

## Abstract

During embryonic development dramatic cell shape changes and movements re-shape the embryonic body plan. These require robust but dynamic linkage between the cell-cell adherens junctions and the force-generating actomyosin cytoskeleton. Our view of this linkage has evolved, and we now realize linkage is mediated by a mechanosensitive multiprotein complex assembled via multivalent connections. Here we combine genetic, cell biological and modeling approaches to define the mechanism of action and functions of an important player, *Drosophila* Polychaetoid, homolog of mammalian ZO-1. Our data reveal that Pyd reinforces cell junctions under elevated tension, and facilitates cell rearrangements. Pyd is important to maintain junctional contractility and in its absence cell rearrangements stall. We next use structured illumination microscopy to define the molecular architecture of cell-cell junctions during these events. The cadherin-catenin complex and Cno both localize to puncta along the junctional membrane, but are differentially enriched in different puncta. Pyd, in contrast, exhibits a distinct localization to strands that extend out from the region occupied by core junction proteins. We then discuss the implications for the protein network at the junction-cytoskeletal interface, suggesting different proteins localize and function in distinct ways but combine to produce robust connections.

## Introduction

One fundamental property of animal cells is their ability to change shape and move. This depends on force generation, with the actomyosin cytoskeleton playing a prominent role. However, for force to be translated into cell shape change and movement, the cytoskeleton must be anchored at the plasma membrane. This occurs at cell-cell and cell matrix junctions, which also serve as signal transduction hubs for sensing and transducing mechanical force and chemical signals (Ladoux and Mege, 2017). One challenge for our field is to define the machinery and mechanisms that link cell-cell adherens junctions (AJ) to the cytoskeleton, allowing tissues to change shape during the complex events of embryonic morphogenesis without disrupting tissue architecture.

Our understanding of the nature of this linkage evolved rapidly over the last two decades. The central role of transmembrane cadherins and their cytoplasmic partners beta- and alpha-catenin in mediating adhesion was defined early on. This initially suggested a simple linear connection to actin, via the ability of alpha-catenin to bind actin. However, new data made this picture increasingly complex, with the addition of many additional proteins at the AJ:cytoskeletal interface and the realization that the structure and function of this multiprotein assemblage is altered by applied force, via mechanosensitive feedback loops (Yap *et al*., 2018; Troyanovsky, 2022). Thus, evolution shaped an exceptionally robust multiprotein machine to accommodate force on AJs during the complex events of morphogenesis (Fernandez-Gonzalez and Peifer, 2022).

We use the fruit fly *Drosophila* as a model, allowing us to combine state-of-the-art genetic and cell biological tools. In *Drosophila*, different proteins occupy distinct places on the spectrum of importance for junctional integrity. The core cadherin-catenin complex and, in most tissues, the polarity protein Bazooka (Baz)/Par3, are essential-their loss disrupts cell adhesion itself (e.g. Cox *et al*., 1996; Müller and Wieschaus, 1996; Tepass *et al*., 1996; Uemura *et al*., 1996; Sarpal *et al*., 2012). Other proteins, like Canoe/Afadin, are essential for many morphogenetic movements and support tissue integrity in tissues under tension, but are not essential for adhesion (Boettner *et al*., 2003; Sawyer *et al*., 2009; Sawyer *et al*., 2011). Finally, some proteins, like Vinculin (Maartens *et al*., 2016), Ajuba (Razzell *et al*., 2018; Rauskolb *et al*., 2019), or Sidekick (Finegan *et al*., 2019; Letizia *et al*., 2019), are not essential for viability or overall morphogenesis, but their loss weakens connections at AJs under tension. In addition, many proteins in the AJ network are complex multidomain proteins—connections linking them are not linear but form a network of multivalent interactions. Thus, one can remove individual domains of important players like Canoe/Afadin (Perez-Vale *et al*., 2021) or even of alpha-catenin (Sheppard *et al*., 2023) without fully disrupting AJ: cytoskeletal connections.

Our current goal is to position Drosophila Polychaetoid (Pyd), the fly homolog of mammalian Zonula occludens (ZO) proteins, into this network and define its cellular and molecular functions. All family members are complex multidomain proteins with three PDZ domains, SH3 and Guanylate kinase (Guk) domains, an actin binding region, and regions of intrinsic disorder. This multivalent structure allows them to interact with a diverse set of protein partners, including transmembrane proteins like claudin, occludin and JAM and adapters like Canoe/Afadin, as well as actin (Rouaud *et al*., 2020). We seek to define roles Pyd plays in the complex events of morphogenesis and in strengthening AJs under tension.

Mammalian ZO-1 and its relatives, ZO-2 and ZO-3 are best known for their role in tight junctions, which seal epithelial sheets and preserve barrier function (Umeda *et al*., 2006). However, they also play a role in AJ establishment in vitro (Itoh *et al*., 1997; Ikenouchi *et al*., 2007). The presence of three mammalian family members has made analyzing their roles in morphogenesis challenging. ZO-3 is dispensable. ZO-2 mutants die during gastrulation (Xu *et al*., 2008), but chimeric embryos in which the embryo proper is almost exclusively derived from ZO-2 mutant cells develop normally to adulthood (Xu *et al*., 2009). ZO-1 mutants gastrulate normally, but exhibit die at E10.5 (Katsuno *et al*., 2008)—however, this might also result from defects in extraembryonic tissues. To define the full function of the ZO family, one would need to generate double or triple mutant embryos. This has not been done, but conditional knockout experiments in the liver (Itoh *et al*., 2021) and kidney (Itoh *et al*., 2018) suggest complete or partial redundancy between ZO-1 and ZO-2.

The presence of a single ZO family member in Drosophila, Pyd, facilitates analysis of its roles. Pyd was identified via alleles that lead to supernumerary adult bristles (Neel, 1940; Chen *et al*., 1996); mutants also have adult eye and embryonic tracheal defects (Jung *et al*., 2006). The gene is complex with 11 potential isoforms and 3 predicted translation start sites, and these original alleles were not null but reduced or altered expression in tissue specific ways. In 2011, two groups generated alleles completely deleting protein coding sequences (Choi *et al*., 2011; Djiane *et al*., 2011). Strikingly, null zygotic mutants were adult viable with defects in bristle number, wing shape and the female germline, revealing that Pyd is not essential for cell-cell adhesion post-embryonic morphogenesis. However, the new alleles allowed the removal of both maternal and zygotic Pyd. This revealed that Pyd, while not essential, is important for high fidelity completion of embryonic morphogenesis. In its absence, 40-60% of embryos die, with defects in the complex movements of head involution and dorsal closure (Choi *et al*., 2011). However, almost half of the *pyd M/Z* null mutants survived embryogenesis. These data suggest Pyd occupies an interesting place in the spectrum of protein importance for AJ robustness. However, these earlier analyses only examined the latest morphogenetic events, focusing on dorsal closure. Thus, our first goal was to define Pyd’s roles during the earlier morphogenetic events when Cno plays important roles. To do so, we assessed localization of other AJ and cytoskeletal proteins, AJ stability, junctional contractility, and cell exchange.

The growing appreciation of the diversity of proteins at the AJ:cytoskeletal interface and their multivalent interactions also raised important questions about the molecular architecture of AJs. AJs are very large molecular assemblies, encompassing hundreds to thousands of copies of E-cadherin and its cytoplasmic binding partners. Scientists studying cell-matrix junctions led the way, suggesting that these junctions have a layered three-dimensional nano-architecture, with different proteins localized to different zones (Case and Waterman, 2015). Analysis of AJ architecture in mammalian epithelial cells using electron microscopy (Efimova and Svitkina, 2018) or elevated resolution light microscopy (e.g. Choi *et al*., 2016; Bertocchi *et al*., 2017) also have begun to support models like this, with different AJ or cytoskeletal proteins along the plasma membrane-proximal to membrane-distal axis. However, how AJ architecture evolves during morphogenesis, and how Pyd fits into the picture remain unanswered questions. Thus, our second goal was to use high resolution microscopy to begin to define the substructure of AJs in vivo during morphogenesis.

## Results

### Pyd reinforces adherens junctions under elevated tension and facilitates cell rearrangements, but it is less essential for junctional stability than Cno

The multiprotein AJ complex includes proteins whose functions range from essential for adhesion, like the core cadherin-catenin complex, to those important for many morphogenetic movements, like Cno, to those that are dispensable for viability and play reinforcing or tissue-specific roles, like Sdk. Our first goal was to place Pyd’s in this network, defining its roles in morphogenesis and comparing them to the roles of Cno.

Unlike *cno*, *pyd* zygotic mutants are viable to adulthood (Seppa *et al*., 2008; Djiane *et al*., 2011). However, most maternal/zygotic (M/Z) *pyd* mutants (40-80% depending on the allelic combination) die as embryos with defects in head involution, dorsal closure and tracheal development (Jung *et al*., 2006; Choi *et al*., 2011), all events occurring near the end of embryonic morphogenesis. During dorsal closure, pyd plays a role in maintaining a straight leading edge and uniform leading edge cells shapes, consistent with a role in reinforcing connections between AJs and the leading edge actomyosin cable. In these phenotypes, M/Z *pyd* mutants resemble zygotic *cno* mutants, which have reduced but not eliminated Cno function, (Choi *et al*., 2011).

Cno plays multiple roles earlier in embryonic development, in events including initial positioning of AJs (Choi *et al*., 2013; Schmidt *et al*., 2018), mesoderm invagination (Sawyer *et al*., 2009), and germband elongation (Sawyer *et al*., 2011). To assess whether Pyd plays roles reinforcing AJ:cytoskeletal connections during these earlier stages, we examined M/Z *pyd* mutants and compared their phenotype to M/Z *cno* mutants. To generate M/Z *pyd* mutant embryos, we crossed homozygous mutant females with heterozygous *pyd* males and collected embryos from this cross. We distinguished M/Z mutants from paternally rescued siblings by staining with Pyd antibody.

One early role for Cno is in apical constriction of cells in the ventral furrow. By stage 8, the ectoderm should have joined at the midline and mesoderm should be fully internalized (Fig. 1A). M/Z *cno* mutants have a fully penetrant defect in this (Sawyer *et al*., 2009). Consistent with Pyd helping reinforce AJs under tension, ventral furrow invagination was defective in many M/Z *pyd* mutants. However, unlike M/Z *cno* mutants, this phenotype was not fully penetrant. Mesoderm invagination went to completion in 24% of stage 7-9 M/Z *pyd* mutants (n= 45 mutant embryos). 53% had mild closure defects at the anterior or posterior ends (Fig. 1A vs B) and only 22% had the severe invagination failure (Fig. 1C) seen in M/Z *cno* mutants. Thus, complete loss of Pyd has less severe effects on mesoderm invagination than loss of Cno—it is also less penetrant in its effects than *cnoΔRA*, which lacks Cno’s Rap1 binding RA domains. Instead Pyd loss is more similar in this phenotype to the milder *cnoΔFAB* mutant, which lacks the C-terminal F-actin binding region (Perez-Vale *et al*., 2021).

**Figure 1.**
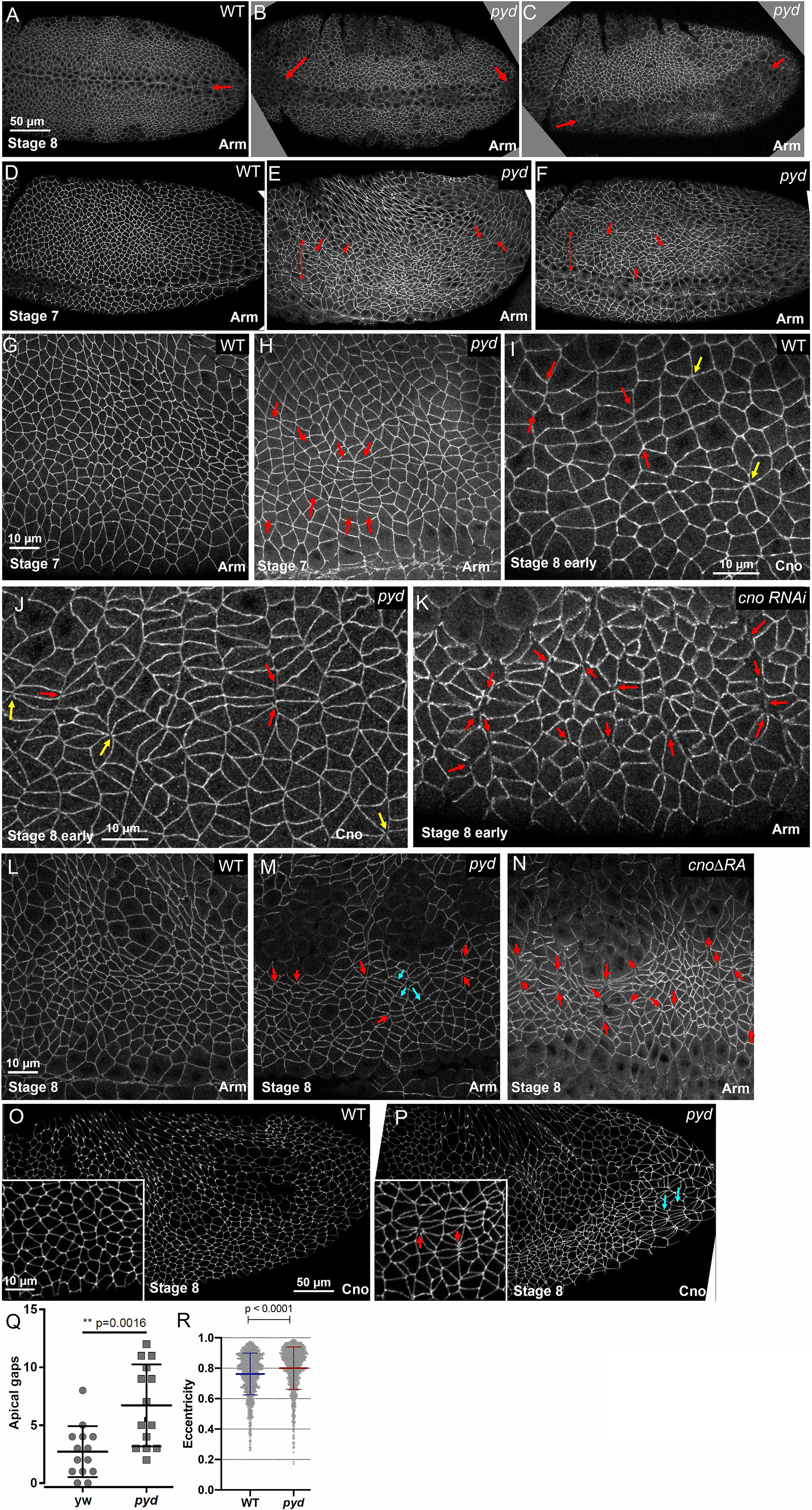
Pyd reinforces adherens junctions under elevated tension and facilitates cell rearrangements, but is less essential for junctional stability than Cno. Unless noted, in this and subsequent Figures embryo images are anterior left and dorsal up. A-C. Ventral views, stage 8 embryos. A. In wildtype the ventral furrow has closed (arrow). B, C. In M/Z *pyd* mutants, most embryos have mild (B, arrow) or more severe (C, arrow) defects in ventral furrow closure. D-H Stage 7 embryos. D,G. Wildtype. E,F,H. In M/Z *pyd* mutants, many cells in the ventral ectoderm (brackets) are more elongated along the anterior posterior axis (arrows). These often form stacks of elongated cells (H, arrows). I-K. Early stage 8. I. In wildtype cell junctions remain intact, including at aligned AP borders (red arrows) or at rosette centers (yellow arrows). J. In M/Z *pyd* mutants occasional apical gaps appear in cell junctions at aligned AP borders (red arrows) or at rosette centers (yellow arrows). K. Gaps are more frequent in embryos in which Cno function is strongly reduced by RNAi. L-P. Stage 8. L vs M. O vs P. In M/Z *pyd* mutants, aligned elongated cells (M,P cyan arrows) and gaps at apical junctions (M, P inset, red arrows) N. Gaps are more frequent and more severe in *cnoΔRA* mutants. Q. Quantification of increased cell eccentricity in M/Z *pyd* mutants. R. Q. Quantification of increased frequency of apical junctional gaps in M/Z *pyd* mutants.

We next looked at germband extension, during which planar polarization of AJ and cytoskeletal proteins drives cell intercalation and body axis extension. In M/Z *cno* mutants and in *cno* mutants lacking its Rap1-binding RA regulatory domains, multiple gaps open at AJs under elevated tension: those at AP borders and tricellular junctions (TCJs; Sawyer *et al*., 2011; Perez-Vale *et al*., 2021). Cell shapes are also altered in *cno* mutants, with cell elongation and preferential cell alignment along the AP axis. We thus examined AJ stability under elevated tension and cell rearrangements during germband elongation in M/Z *pyd* mutants. We focused on the thoracic and abdominal region during germband extension, which starts in embryonic stage 7 when cells began intercalating.

Two defects were noted in M/Z *pyd* mutants. First, occasional apical gaps or more subtle disruptions were observed at TCJs/rosettes or aligned AP borders (Fig. 1I vs J, yellow or red arrows, respectively). Quantification verified an increase in gaps at stages 7-8 (Fig. 1Q). However, the gaps were not as numerous as those seen after *cno* RNAi (Fig. 1K, red arrows), or in embryos mutant for *cnoΔRA* (Fig. 1N red arrows; (Perez-Vale *et al*., 2021). Instead, the severity of the gap phenotype in M/Z *pyd* mutants was more similar to that seen in the milder *cnoΔFAB* mutant (Perez-Vale *et al*., 2021). Thus, Pyd plays a role in stabilizing AJ-cytoskeletal connections under tension, but its role is more modest than that of Cno. Second, in M/Z *pyd* mutants apical cell shapes became more variable (Fig. 1D vs E,F), with the appearance of stacks of cells in the lateral ectoderm that were more elongated along the AP axis (Fig. 1E-F arrows, G vs H arrows). Quantification of cell eccentricity confirmed this cell elongation (Fig. 1R). This alteration in cell shape resembled what we saw in *cnoΔRA* mutants (Perez-Vale *et al*., 2021). These defects continued at stage 8 in M/Z *pyd* mutants, with continued presence of stacks of elongated cells (Fig. 1L vs M, cyan arrows) and an elevated number of gaps (Fig. 1M, red arrows; Fig. 1O vs P, insets). Thus, by both measures, the phenotypes of M/Z *pyd* mutants were similar to but quantitatively less severe than those of M/Z *cno* mutants.

### Pyd helps restrain planar polarity of a subset of AJ proteins, but its loss does not lead to obvious detachment of myosin or actin from AP cell borders

Germband extension is driven in part by reciprocal planar polarization of the actomyosin cytoskeleton to AP borders and junctional proteins to DV borders. Myosin and F-actin enriched on anterior-posterior (AP) cell borders power myosin-driven contractility and thus T1 cell transitions and formation of multicellular rosettes. Cadherin-catenin complex proteins and especially Bazooka (Baz)/Par3 are enriched on dorsal-ventral (DV) cell borders (Fig. 2A, C” cyan vs yellow arrows; Bertet *et al*., 2004; Zallen and Wieschaus, 2004). Intriguingly, Cno is enriched on AP borders while Pyd is enriched at DV borders raising the possibility that each reinforces distinct regions (Manning *et al*., 2019). Cno is also enriched at TCJs, which are sites of elevated junctional tension as AP borders constrict (Sawyer *et al*., 2009; Yu and Zallen, 2020). Cno loss leads to detachment of myosin cables from AP borders. Arm, Pyd, and especially Baz are reduced on AP borders, strongly elevating their planar polarity (Sawyer *et al*., 2011; Manning *et al*., 2019). However, myosin planar polarity is not altered (Sawyer *et al*., 2011).

**Figure 2.**
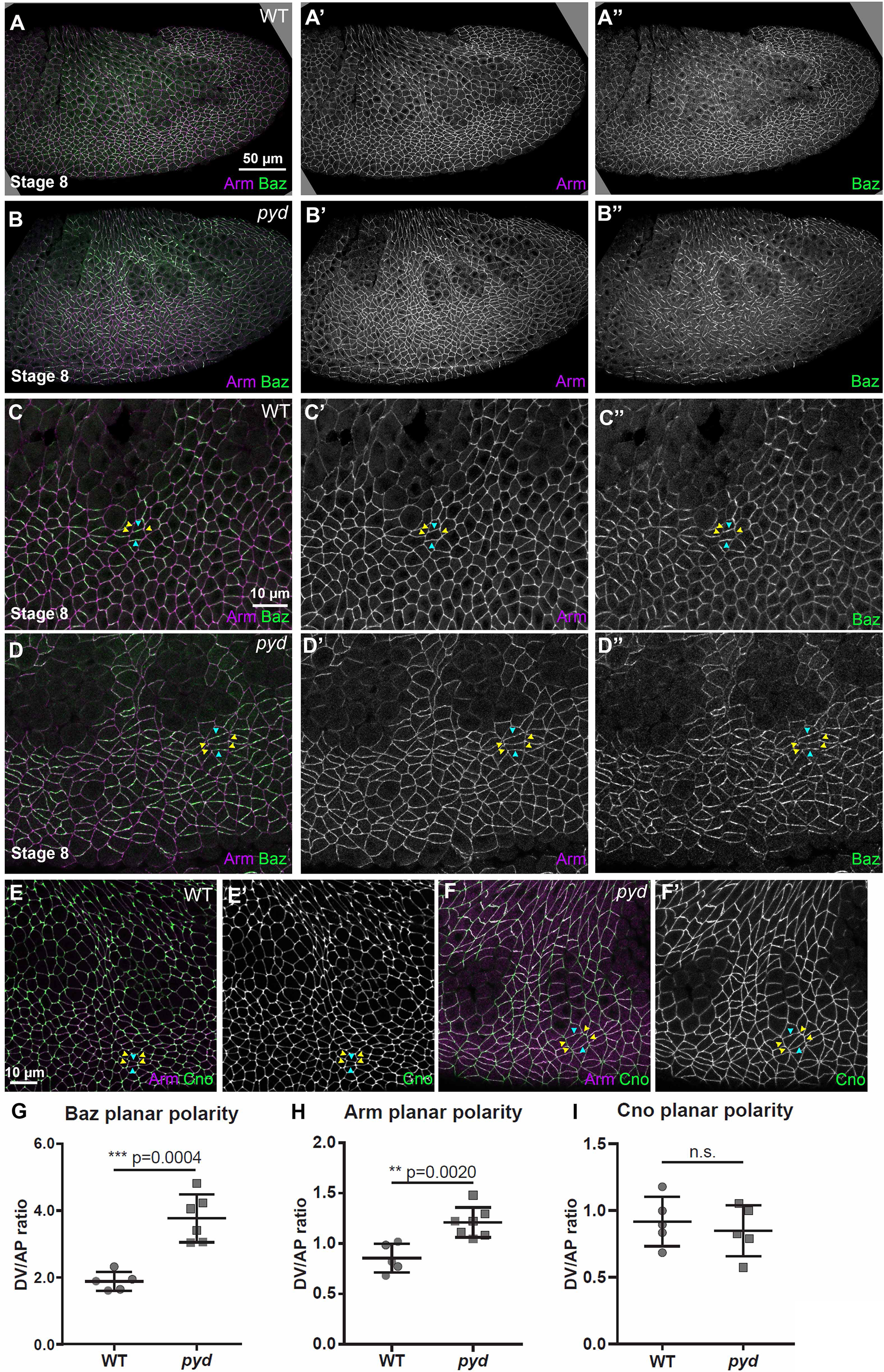
Pyd is required to restrain planar polarity of a subset of AJ proteins. Stage 8 embryos. A-D. Arm and Baz. A,C. In wildtype Arm is relatively uniform around the cells, while Baz is enriched on DV vs AP borders (cyan vs. yellow arrowheads). B,D. In M/Z *pyd* mutants Arm is subtly reduced at AP borders (yellow arrowheads) and Baz is also reduced, thus enhancing the planar polarity of both. E. In wildtype Cno is subtly enriched on AP vs DV borders (yellow vs cyan arrowheads). F. Cno planar polarity is not substantially altered in M/Z *pyd* mutants. G-I. Quantification of Baz, Arm and Cno planar polarity in wildtype and M/Z *pyd* mutants. Error bars represent SD, the significance was determined by an unpaired two-tailed t-test.

We thus examined whether Pyd is important for normal junctional planar polarity. In M/Z *pyd* mutants Baz planar polarity is strongly enhanced (Fig 2A vs B; C vs D, cyan vs yellow arrows; quantified in Fig 2G), and Arm planar polarity is also subtly elevated (Fig 2C” vs D”; quantified in Fig 2H). However, in M/Z *pyd* mutants Cno remained somewhat enriched on AP borders (Fig. 2E vs F; quantified in 2I), contrasting with the reversal of Cno planar polarity seen after loss of its regulator Rap1 (Perez-Vale *et al*., 2023) or deletion of Cno’s RA domain (Perez-Vale *et al*., 2021). Thus, Pyd restrains planar polarity of some but not all AJ proteins. We also examined whether the tight connection of myosin to AJs was disrupted. In wildtype myosin is planar polarized to AP borders during germband extension, and tightly apposed there (Fig. 3A, D arrows). In M/Z *pyd* mutants, other than places where occasional junction gaps were observed, myosin remained planar polarized and tightly apposed to AP borders (Fig. 3B, E arrows). This contrasted with the clear detachment of myosin from AP borders seen in M/Z *cno* mutants (Fig. 3C; Sawyer *et al*., 2011). We also examined F-actin localization. Once again, F-actin did not appear to detach from AP cell borders in M/Z *pyd* mutants (Fig. 3F, cyan arrows), except where junctional gaps appeared (Fig. 3F, yellow arrows). This contrasts with M/Z *cno* mutants (Sawyer *et al*., 2011). Together these data suggest Pyd regulates AJ protein planar polarity, but reveal that its role is not as essential as that of Cno. They also reveal that Pyd loss does not lead to myosin detachment from AP borders.

**Figure 3.**
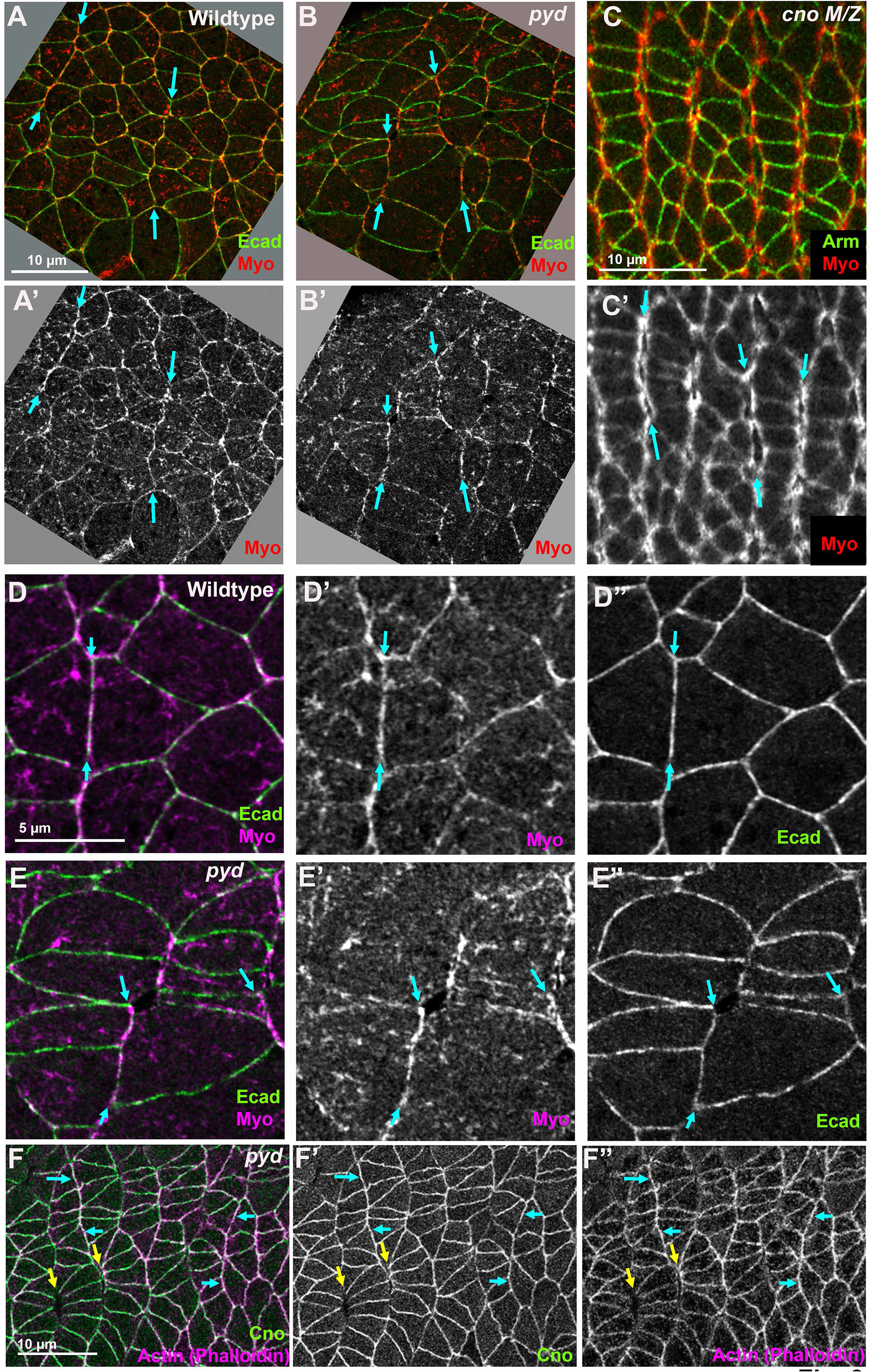
Loss of Pyd does not lead to obvious detachment of myosin from AP cell borders, in contrast with loss of Cno. A-E. Stage 7 embryos, expressing Ecad-GFP and mCh-Myosin. A,D. In wildtype, junctional myosin is enriched at AP borders (arrows), where it is tightly apposed to junctional Ecad. B,E. This remains true in M/Z *pyd* mutants (arrows), except where there are junctional gaps. C. In contrast, in M/Z *cno^R2^* null mutants, Myosin often detaches from junctions at AP borders (arrows). F. F-actin, as revealed using Phalloidin, does not detach from AP borders in M/Z *pyd* mutants (cyan arrows), except places where junctions have detached as marked by Cno (yellow arrows).

### Pyd and Cno are mutually required for one another’s enrichment at TCJs

When AJs are put under molecular tension, their connection to the cytoskeleton is strengthened by conformational changes and protein recruitment. For example, Cno is recruited to AJs under tension, including TCJs during germband extension (Fig 4A, arrows; 4C; Sawyer *et al*., 2009; Yu and Zallen, 2020), where it is important for TCJ stabilization (Sawyer *et al*., 2009; Perez-Vale *et al*., 2021). Pyd is enriched at TCJs in larval wing imaginal discs and was reported to be enriched at TCJs in stage 9-11 embryos (Letizia *et al*., 2019). Consistent with this, during stage 7 Pyd is modestly enriched at TCJs (Fig. 4D, E; ∼1.5 fold). Pyd enrichment at TCJs increases as development proceeds, with ∼2-fold enrichment at stages 9-10 (Fig. S1A, C).

**Figure 4.**
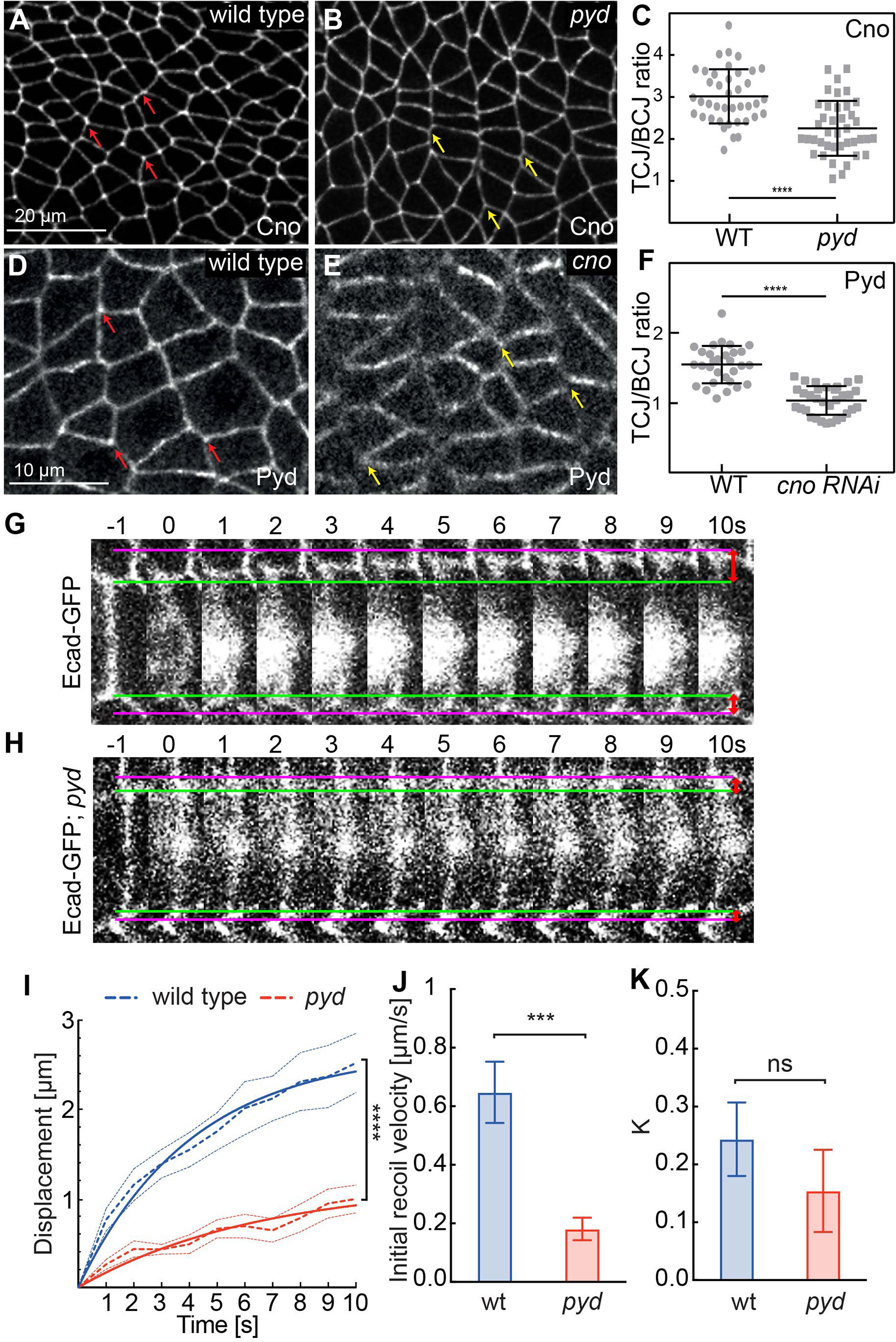
Pyd and Cno are mutually required for one another’s enrichment at TCJs, and M/Z *pyd* mutants have reduced junctional tension. A,B,D,E. Stage 7 embryos. A. In wildtype Cno is enriched at TCJs (arrows). B. Cno TCJ enrichment is reduced in M/Z *pyd* mutants (arrows). C. Quantification of Cno TCJ enrichment. D. In wildtype Pyd is mildly enriched at TCJs (arrows). E. *cno* RNAi reduces Pyd junctional localization and TCJ enrichment (arrows). F. Quantification of Pyd TCJ enrichment. G, H. Still frames from movies of Stage 7 embryos, expressing Ecad-GFP. Junctions were severed with a laser and junctional recoil was measured. Green line=TCJs before cutting. Purple line=TCJ position after 10 s. I. Vertex displacement versus time plot for junction cuts. Dashed curves represent the mean with SEM (error bars) of displacement of junction cuts from the wild type and *pyd* mutant embryos during germband extension. Solid curves represent the one-phase association fitting of experimental data. N=10 junctions from 8 wild type embryos and 10 junctions from 7 *pyd* mutant embryos, two-way ANOVA, Sidak’s multiple comparisons test, **** p<0.0001. J Mean with SEM of initial recoil velocities from the one-phase association fitting in (I). Unpaired two-sided t-test estimate the p value, *** p<0.001. K. Mean with SEM of K values from the one-phase association fitting in (I), which indicate the ratio between the elasticity of the junction and the viscosity coefficient. Unpaired two-sided t-test estimate the p value, ns, not significant.

The TCJ-enriched protein Sidekick is important for full TCJ enrichment of Cno in stage 9-11 embryos (Letizia *et al*., 2019). We thus asked whether Pyd is also required for Cno TCJ enrichment. In M/Z *pyd* mutants, Cno TCJ enrichment at stage 7 was significantly reduced, but not eliminated (Fig. 4A vs B arrows; quantified in 4C). Sidekick is also important for Pyd enrichment in imaginal discs (Letizia *et al*., 2019). We thus wondered whether Cno was similarly required for Pyd TCJ enrichment. We used a validated strong *cno* RNAi line to knockdown Cno maternally and zygotically (Bonello *et al*., 2018; Manning *et al*., 2019), and examined effects on Pyd TCJ enrichment. To confirm Cno knockdown, we co-stained the embryos with Cno antibody. Low levels of Cno were still detectable (Fig S1B). Cno knockdown led to gaps at AJs under tension, as previously noted (Manning *et al*., 2019), and reduced uniform Pyd recruitment at AJs. At stage 8 Pyd TCJ enrichment was largely abolished after *cno* RNAi (Fig. 4D arrows; quantified in 4G), and this continued at stage 9/10 (Fig. S1B,C). These data are consistent with Pyd being recruited to TCJs under elevated tension, and suggests that the multiple proteins that are enriched at TCJs mutually reinforce one another’s recruitment.

### M/Z *pyd* mutants have reduced junctional tension

The reduction in Cno TCJ recruitment in M/Z *pyd* mutants and the hyper planar polarization of Baz to DV borders made us wonder whether loss of Pyd might alter junctional tension. To test this, we used laser cutting to sever the apical junctional cortex in stage **7** embryos, and measured displacement of the TCJs flanking the severed AJ. In wildtype embryos there is a rapid recoil of ∼2.4µm within 10 seconds (Fig. 4G,I), with a mean initial recoil velocity of ∼0.6 µm/sec (Fig. 4J). These values were reduced in M/Z *pyd* mutants—thee was a total recoil of ∼1µm in 10 seconds (Fig. 4H,I) with a mean initial recoil velocity of ∼0.2 µm/sec (Fig 4J). There was no significant change in the ratio between the elasticity of the junction and the viscosity coefficient (k; Fig. 4K), supporting the idea that the tension on the junctions is affected. Thus, loss of Pyd reduces junctional tension, consistent with the reduction of Cno TCJ recruitment.

### Modeling and experimental analysis suggest T1 transitions are slowed or fail in M/Z *pyd* mutants

One of the most striking features of M/Z *pyd* mutants was the appearance of stacks of cells elongated along the AP axis. Similar cell shape defects were observed during germband extension after the loss of other AJ:cytoskeletal linkers (e.g., Sawyer *et al*., 2011; Tamada *et al*., 2012; Razzell *et al*., 2018; Finegan *et al*., 2019). Planar-polarized myosin contractility at AP borders drives cells into the T1 configuration or into rosettes, which then resolve to extend the DV axis. Our data above reveal that loss of Pyd reduces junctional contractility. We thus wondered whether reducing border contractility could lead to the stacks of cells elongated along the AP axis we observed in M/Z *pyd* mutants.

To test this, we turned to a vertex model of germband extension created to explore cell interface behaviors of the germband during axis extension (Tetley *et al*., 2016). We adapted an updated version of this germband extension vertex model that includes the extrinsic posterior pulling force of the midgut (Finegan *et al*., 2019), to test whether reduced contractility recapitulated the cell stacks elongated along the AP axis. We ran the simulation for 400 timesteps, equating to roughly 20 simulated minutes of germband extension (∼0.5 min of germband extension per timestep). Starting configurations were the same, and we examined cell shapes and intercalation at 10 minutes and 20 minutes of simulated germband extension. In the wildtype, cells effectively intercalated and the germband extended, with cells retaining largely isotropic shapes (Fig. 5A; Suppl. Movie 1). In contrast, when ‘supercontractilty’, modelled by a condition where junction shrinkage rate increases as length decreases, was removed to mimic the reduction in contractility seen in pyd mutant, cell intercalation was reduced, and the cell shape and arrangement phenotype, with stacks of cells elongated along the AP axis, matched that we see in M/Z pyd mutants (Fig. 5B; Suppl. Movie 2).

**Figure 5:**
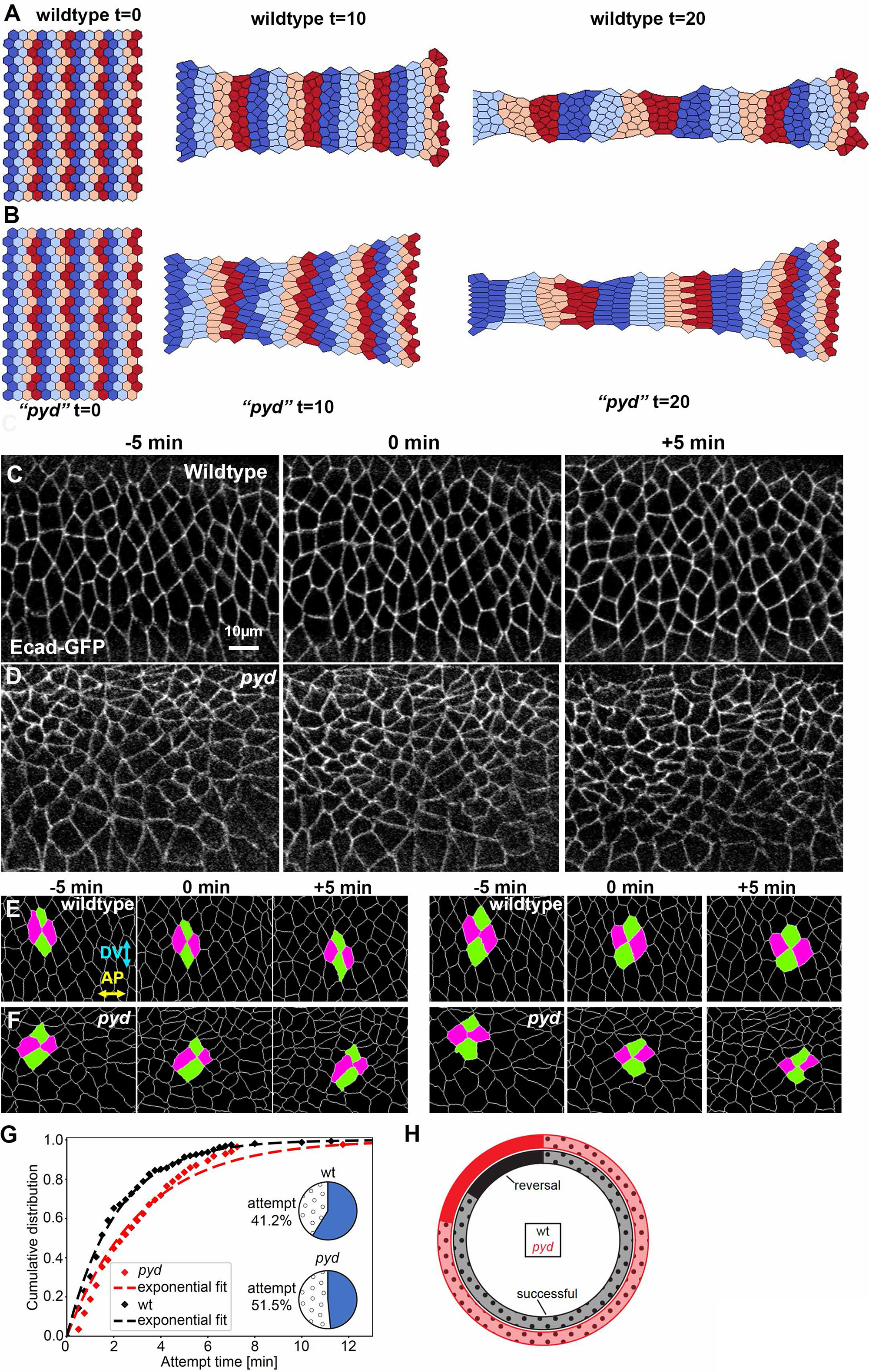
Modeling suggests reducing contractility can mimic *pyd* phenotypes, while experiments reveal that T1 transitions in *pyd* M/Z embryos are slowed and reverse more often. A, B. Stills from three time points in the vertex model simulations of wildtype germband extension and germband extension with reduced contractility, to model *pyd* mutants. Distinct colors correspond to cells in different parasegments. C-D. Stills of movies from wildtype (C) and *pyd* M/Z (D) embryos expressing ECadherin-GFP. E,F. Examples of T1 transitions of wildtype (E) and *pyd* (D) embryos are displayed. The old neighbors are marked in magenta while new neighbors are in green. Two successful T1 transitions are shown for wildtype (green frames) while for *pyd* mutants one successful (green frame) and one unsuccessful (red frame) transition is displayed. G. The cumulative distribution of attempted T1s in wildtype (black) and *pyd* mutant (red) embryos was plotted against time. The segmented lines display exponentially fitted curves to extract half times, indicating that *pyd* mutant embryos needed more time to finish a T1 attempt. The pie graphs display the percentage of direct (blue) and attempted (circles) T1s in wildtype and *pyd* mutant embryos. (F) In wildtype, 84% of T1 transitions elongate the newly formed border in AP (horizontal) direction, while 16% fail and reverse after formation of a 4x vertex in wildtype (black doughnut; n=327 transitions). The number of reversed T1 transitions was elevated to 22% in *pyd* mutants (red doughnut; n=216 transitions).

This modeling suggested the possibility that the reduced AJ contractility we saw in M/Z *pyd* mutants might affect T1 transitions, in which four cells constrict their AP (vertical) borders to join at a single vertex, and they exit by elongating their DV (horizontal) junctions (Fig. 5E). To determine if M/Z *pyd* mutants have a defect in T1 transitions, we live imaged embryos in stage 7-8 expressing endogenously tagged E-Cadherin GFP (Fig. 5C,D). Movies were then segmented to define cell junctions, and the rate and direction of T1 transitions were analyzed. We found that in wildtype embryos the majority of T1 transitions went directly from the formation of a 4 cell vertex into the elongation of the new junction along the AP axis (Fig. 5E). Those T1 transitions were scored as direct transitions without an attempt phase. In contrast, when an extending 4 cell vertex reversed and returned to a 4 cell vertex, only to once again elongate a junction (without defining the direction in which a new junction was being formed), we scored this as an “attempted T1 transition”. In wildtype embryos 58.4% of T1 transitions proceeded directly while 41.6% were scored as attempted T1 transitions (Fig. 5G). In contrast, in M/Z *pyd* mutant embryos, only 47.7% of T1 transitions proceeded directly, while 52.3% were scored as attempted T1 transitions (Fig. 5D,G). We then calculated the duration of the attempt phase in wildtype versus M/Z *pyd* mutant embryos. The mean attempt duration in wildtype embryos was 2.1 minutes while the mean attempt duration was increased in M/Z *pyd* mutants to 3.2 minutes (Fig. 5G). Finally, we scored the fraction of successful T1 transitions. In wildtype embryos, 84% of formed 4 cell vertices successfully resulted in a horizontal (AP) elongation of the new border (Fig. 5F, H; n=551 transitions). However, in *pyd* mutants, only 78% of T1 transitions were successful, while 22% of 4x vertexes reverted back (Fig. 5F, H). Together, these data are consistent with the idea that slowed and failed T1 transitions can help explain the stacks of elongated cells we observed in M/Z *pyd* mutant embryos.

Putting this together, our phenotypic data suggest that Pyd reinforces TCJs under elevated tension and restrains hyper-planar polarization of some AJ proteins, but reveal that junctional disruptions are relatively rare in its absence. Cell shapes are altered, with stacks of elongated cells accumulating. Pyd loss reduces but does not eliminate junctional tension, and in doing so reduces TCJ enrichment of Cno. Our modeling suggests that this reduction in junctional tension could impede shrinking of AP borders and intercalation, and our analysis of T1 transitions is consistent with this. This positions Pyd in a new place on the continuum of AJ proteins—less essential than Cno, but more essential than proteins like Sdk or Ajuba, where null mutants are viable and fertile. It also provides mechanistic insights into how Pyd loss alters germband extension.

### High resolution microscopy reveals differential enrichment of the cadherin-catenin complex and Cno in junctional puncta, suggesting AJs have substructure

This continuum of functional importance provided evidence that connections between the cadherin-catenin complex and the cytoskeleton are not the simple, linear connections initially envisaged. We wondered how the supermolecular AJ protein network assembles to create this complex and robust connection. When AJs assemble during cellularization, they form large puncta known as spot AJs (Tepass and Hartenstein, 1994), each of which contains ∼1500 cadherin-catenin complexes (McGill *et al*., 2009). At that stage there are roughly 3-8 puncta along each roughly 3-5 µm BCJ (e.g. Harris and Peifer, 2004; McGill *et al*., 2009). As gastrulation begins, the punctate character of AJs becomes more complex, and standard confocal microscopy makes junctions look more continuous. To reveal underlying complexity, we used two forms of high-resolution microscopy, structured illumination (SIM) and the Zeiss Airyscan module to examine AJs at higher lateral resolution, theoretically reaching 100 (SIM)–120nm (Airyscan).

Both approaches revealed the punctate nature of the core cadherin-catenin complex. As the germband elongates, cell shapes become much less regular, with longer and shorter bicellular borders. At stage 8, SIM revealed small Arm puncta that are tightly aligned along the plasma membrane along bicellular junctions (BCJs; Fig 6A). We saw a similar distribution of small puncta aligned along BCJs when imaging Ecadherin-GFP using the Zeiss Airyscan detector (Fig. 6B), although they were not quite as well resolved. We used the Imaris Spot function to analyze our SIM images. This suggested there are 20-50 Arm puncta per 10µm of BCJ (Fig. 6C). Average puncta size was 0.24 µm, with the extremes of puncta measured extending from 0.15 µm-0.4 µm. We next used SIM to explore Arm localization relative to that of Cno. Cno was enriched at AP cell borders (vertical in these images). Cno localized to small junctional puncta along BCJs (Fig 6D”,E”). While the Arm and Cno puncta sometimes overlapped, many puncta were differentially enriched for either Arm or Cno (Fig. 6E-E”, red vs green arrows). Cno was strongly enriched at TCJs (Fig 6F). To discern relative localization of Arm and Cno at TCJs, we turned down the Cno signal (Fig. 6F’). Cno strongly localized to one or more puncta in the center of the TCJ (Fig. 6F’,F’” green arrow; 6G), with Arm more enriched in puncta around the TCJ periphery (Fig. 6F’,F” red arrows, 6G). Processing the images with Imaris allowed us to visualize the BCJ puncta along the X-Z plane, confirming differential enrichment of Arm and Cno in different puncta (Fig. 6H). Finally, we measured the breadth of the signal perpendicular to the junctional plasma membrane. The Cno signal spanned a slightly broader region than that of Arm (Fig. 6I), consistent with Arm being directly bound to the cadherin tail and Cno associated by multivalent interactions with actin and other junctional proteins Thus, while Arm and Cno both localize to AJs, they are differentially enriched in different puncta.

**Figure 6.**
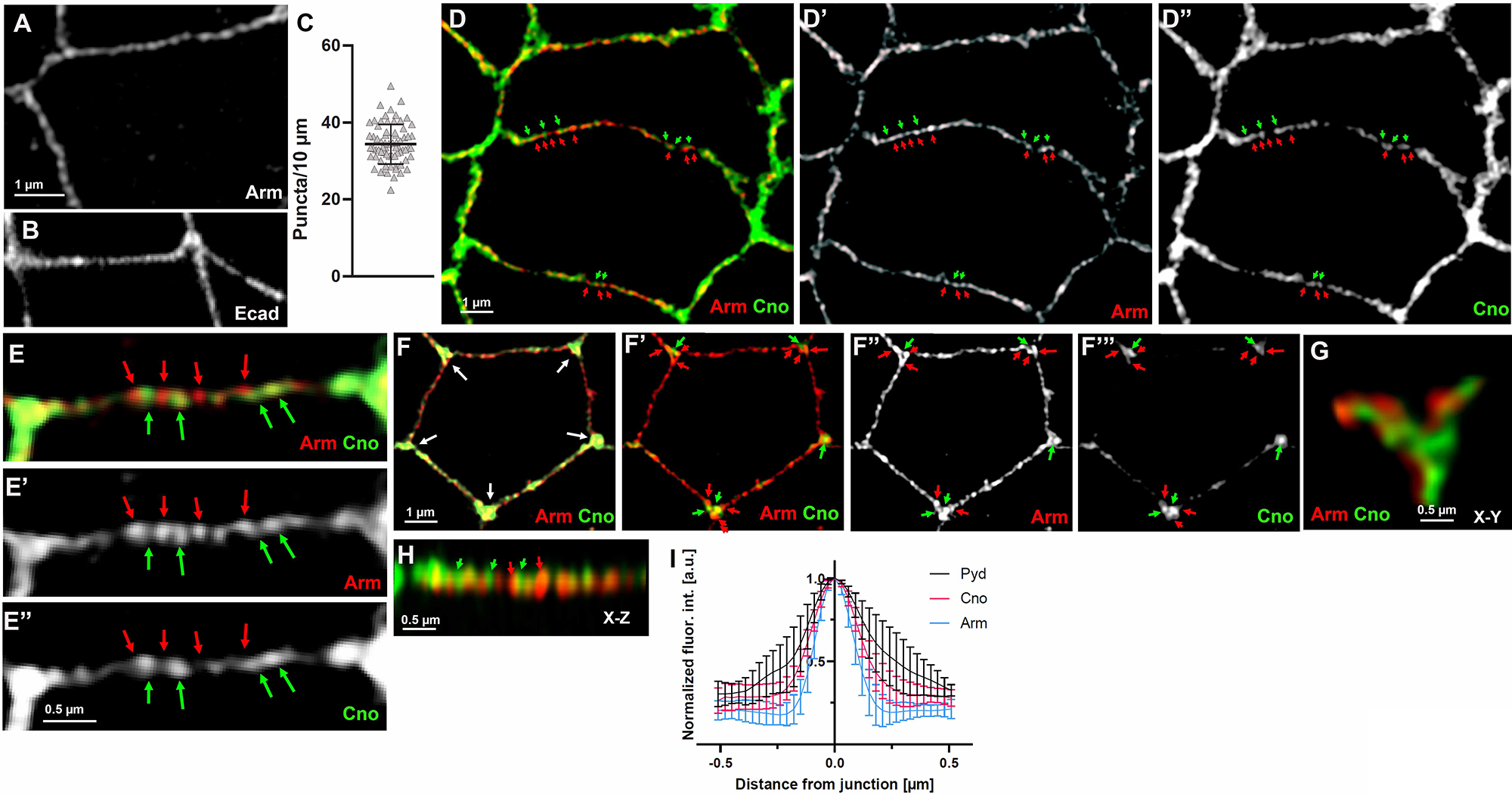
The cadherin-catenin complex and Cno are differentially in junctional puncta along bicellular borders and at TCJs. A, D-H. SIM imaging, stage 7-8 embryos. A. Arm localizes to small puncta along bicellular borders. B. Similar puncta are seen when visualizing Ecad-GFP using the Airyscan module. C. Quantification of Arm puncta per 10 µm. D,E. Cno is enriched at AP borders and TCJs. Cno and Arm both localize to puncta along bicellular borders, but some puncta are more enriched for Arm (red arrows) and others are more enriched for Cno (green arrows). F. Cno is strongly enriched at TCJs (F, white arrows). When the Cno signal is artificially reduced (F’), it reveals that Arm and Cno (red vs green arrows) are enriched in different puncta at TCJs. G,H. Images of representative TCJ and BCJ 3D rendered in Imaris—G is in X-Y plane and H in the X-Z plane. G. Differential localization of Arm and Cno at a TCJ, viewed in the X-Y plane. H. Differential localization of Arm and Cno at a bicellular border, viewed in the X-Z plane. I. Breadth of signal of Arm, Cno, and Pyd, assessed perpendicular to the plasma membranes. Error bars represent SD.

### Pyd localizes to strands that span a broader region than the core junctional proteins

We next used SIM imaging to examine Pyd localization relative to that of Arm or Cno. Here the result was even more striking. In using our Pyd antisera in standard confocal imaging, we had been puzzled that its localization was not as “sharply defined” as that of Arm or Cno (e.g., Fig. 7A vs A’). SIM imaging revealed the reason. Rather than tightly localizing to puncta along bicellular borders, Pyd localized to “strands” over a broader area (Fig. 7B, C). Some overlapped the tighter localization of Arm, but projected membrane distal (Fig. 7B,C, cyan arrows). Other less robust strands projected from the core AJ even further into the membrane-distal zone (Fig. 7B,C yellow arrows). Because this localization was so surprising, we also imaged Arm and Pyd using the Airyscan module, offering a different way to elevate resolution. This also revealed a broader distribution of Pyd, with strands extending membrane-distal from core AJ proteins (Fig. 7D). Quantifying the span of the Pyd signal supported the idea that it extends more plasma membrane distal than Arm or Cno (Fig. 6I). Thus, Pyd occupies a region of the AJ complex quite different from the more core AJ proteins.

**Figure 7.**
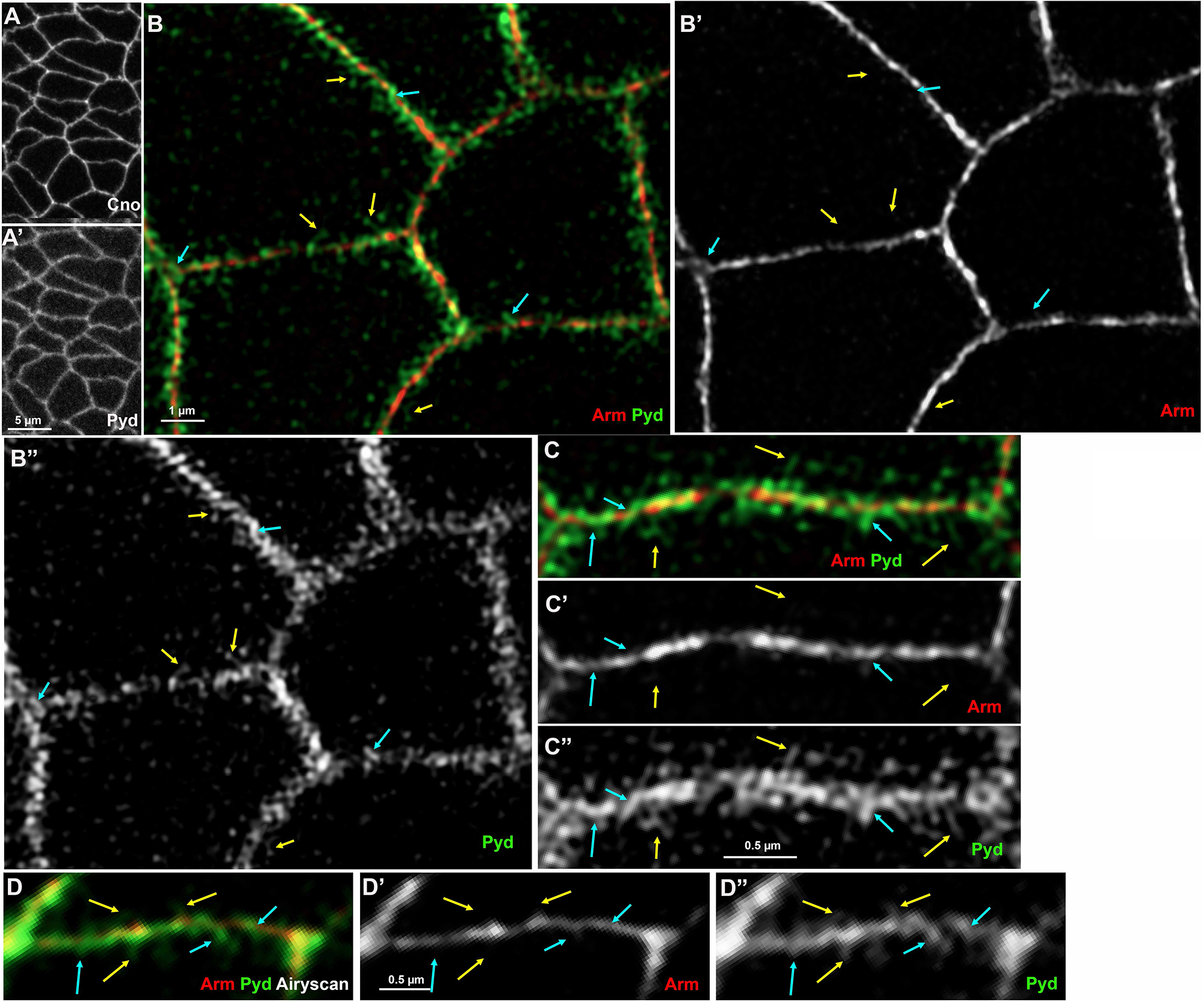
Pyd localizes to strands that span a broader region than the core junctional proteins. A. Standard confocal imaging. Pyd signal appears less tightly localized to junctions than Cno. B-C. SIM imaging, stage 8 embryos. While Arm localizes tightly to puncta along the plasma membranes, Pyd localization extends more broadly. Pyd localizes to strands that align along or perpendicular to the AJs (cyan arrows), and also can be seen localized to finer strands that project into the cytoplasm (yellow arrows). D. Airyscan imaging. While images are less sharp, Pyd localization is also revealed to be much broader than that of Arm.

### Baz and Arm are differentially enriched in different junctional puncta

Next, we used Airyscan imaging to compare the localization of Arm with that of the apical junctional and polarity protein Bazooka (fly Par). While Baz co-localizes with the cadherin-catenin complex in the spot AJs during cellularization (Harris and Peifer, 2004), Baz then becomes successively more apically enriched (Harris and Peifer, 2005). In stage 7 embryos this was apparent in our z-stacks, as the Baz signal was considerably brighter apically, while the brightest signal Arm signal was 0.38µm more basal (Fig. 8A vs B; brightness not adjusted). When examined in a single slice, Arm and Baz localizations overlapped, but Arm and Baz were differentially enriched in different puncta along BCJs (Fig. 8C, red vs green arrows). Quantification perpendicular to the membrane was consistent with the Baz zone being a bit broader than that of the cadherin-catenin complex, but not as broad as the distribution of Pyd along the same junctions (Fig. 8D). Thus, Baz and Arm localize differentially along both the apical-basal axis and along the X-Y plane of bicellular junctions.

**Figure 8.**
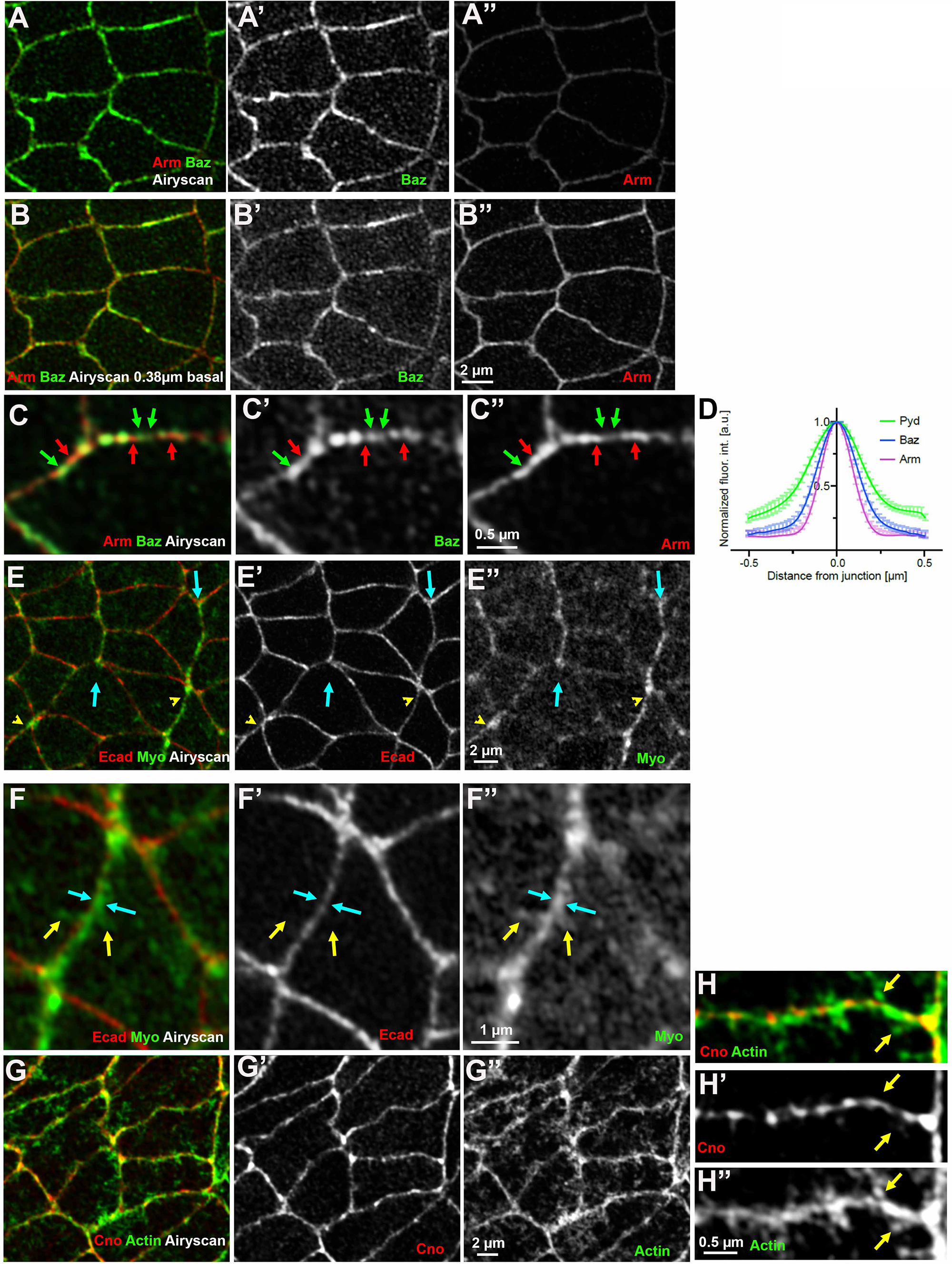
Baz and Arm are differentially enriched in junctional puncta, while Myosin occupies a broader zone along bicellular junctions than core AJ proteins. A-C, E-H, Stages 7-8, Airyscan imaging. A,B. Arm and Baz in Z-sections 0.39µm apart. Baz is stronger in the more apical section, while Arm is stronger in the more basal section. C. Arm and Baz overlap but are differentially enriched in puncta along bicellular borders—some puncta are enriched for Arm (red arrows) and some for Baz (green arrows). D. Breadth of signal of Arm, Baz, and Pyd, assessed perpendicular to the plasma membranes. Error bars represent SD, the significance was determined by an unpaired two-tailed t-test. E,F. Embryo expressing Ecad-GFP and mCh-Myosin. Myosin is enriched at AP borders (D, arrows) and rosette centers (D, arrowheads). Myosin localization at these borders is broader than that of Arm (E, cyan arrows) and strands extend membrane distal (E, yellow arrows). G, H. Actin localization at cell junctions is broader than that of Cno. H. Arrows indicate actin structures extending membrane distal.

### Myosin and actin occupy a broader zone along the bicellular junctions than the core AJ proteins

We also were curious about the localization of the cytoskeleton relative to that of the core junctional proteins. During germband elongation, myosin is present in two broad pools—a pool in the apical membrane undergoing waves of periodic condensation, contraction, and dissipation (Rauzi *et al*., 2010; Fernandez-Gonzalez and Zallen, 2011; Sawyer *et al*., 2011), and a contractile pool at the AJs (Bertet *et al*., 2004; Zallen and Wieschaus, 2004). The AJ pool is planar polarized, with enrichment at AP borders (Fig. 8E, cyan arrows), and at multicellular junctions at the center of rosettes (Fig. 8E, yellow arrowheads; (Bertet *et al*., 2004; Zallen and Wieschaus, 2004). We used Airyscan imaging to compare the localization of Ecad and Myosin. Ecad localized to puncta along each bicellular border (Fig. 8F, F’). Myosin overlapped Ecad but localized to puncta and strands occupying a broader zone that extended membrane distal from Ecad (Fig. 8F, F”). These data are broadly consistent with lovely EM work done in cultured endothelial cells (Efimova and Svitkina, 2018), which also revealed myosin localized membrane distal to the cadherin-catenin complex. We also used phalloidin to visualize F-actin. Similar to myosin, actin localization extended in strands further into the cytoplasm than Cno (Fig. 8G, H arrows). Thus, the cytoskeletal zone is broader than that of the core junctional proteins.

### Using SIM imaging to explore the role of Pyd in AJ molecular architecture

In our final set of experiments, we used SIM imaging to compare AJ architecture in wildtype and in M/Z *pyd* mutants. In wildtype Arm and Cno localize in overlapping but not identical ways to junctional puncta along bicellular borders (Fig. 9A, B arrows). This remained essentially unchanged in M/Z *pyd* mutants (Fig. 9C, D arrows), except in regions where gaps appeared in junctions (Fig. 9C, arrowhead). Arm puncta density along bicellular junctions remained similar (Fig. 9I) and Arm puncta remained tightly apposed to the membrane (Fig 9J). We also examined the effect of loss of Pyd on Baz localization. The overlapping but occasionally differential localization of Arm and Baz to puncta along the bicellular junctions was not drastically altered (Fig. 9E,F vs G,H). However, quantification revealed that the distribution of Baz was somewhat broader in localization to M/Z *pyd* mutants relative to wildtype (Fig. 9K). As a final test of potential roles for Pyd in AJ architecture, we examined whether it affected mobility of Ecad as assessed by Fluorescence recovery after photobleaching (FRAP). We bleached a region of the AJ in embryos expressing Ecad-GFP, and quantified fluorescence recovery (Fig. 9M). In wildtype, recovery was roughly similar to what others have observed (Cliffe *et al*., 2004; Greig and Bulgakova, 2021)—recovery plateaued after 400-500 seconds and the mobile fraction was about 75%. We then repeated this analysis in M/Z *pyd* mutants. No change was seen in either rate of recovery or mobile fraction (Fig. 9N). Thus, Pyd is not essential for many aspects of AJ molecular architecture, but may have effects on Baz localization.

**Figure 9.**
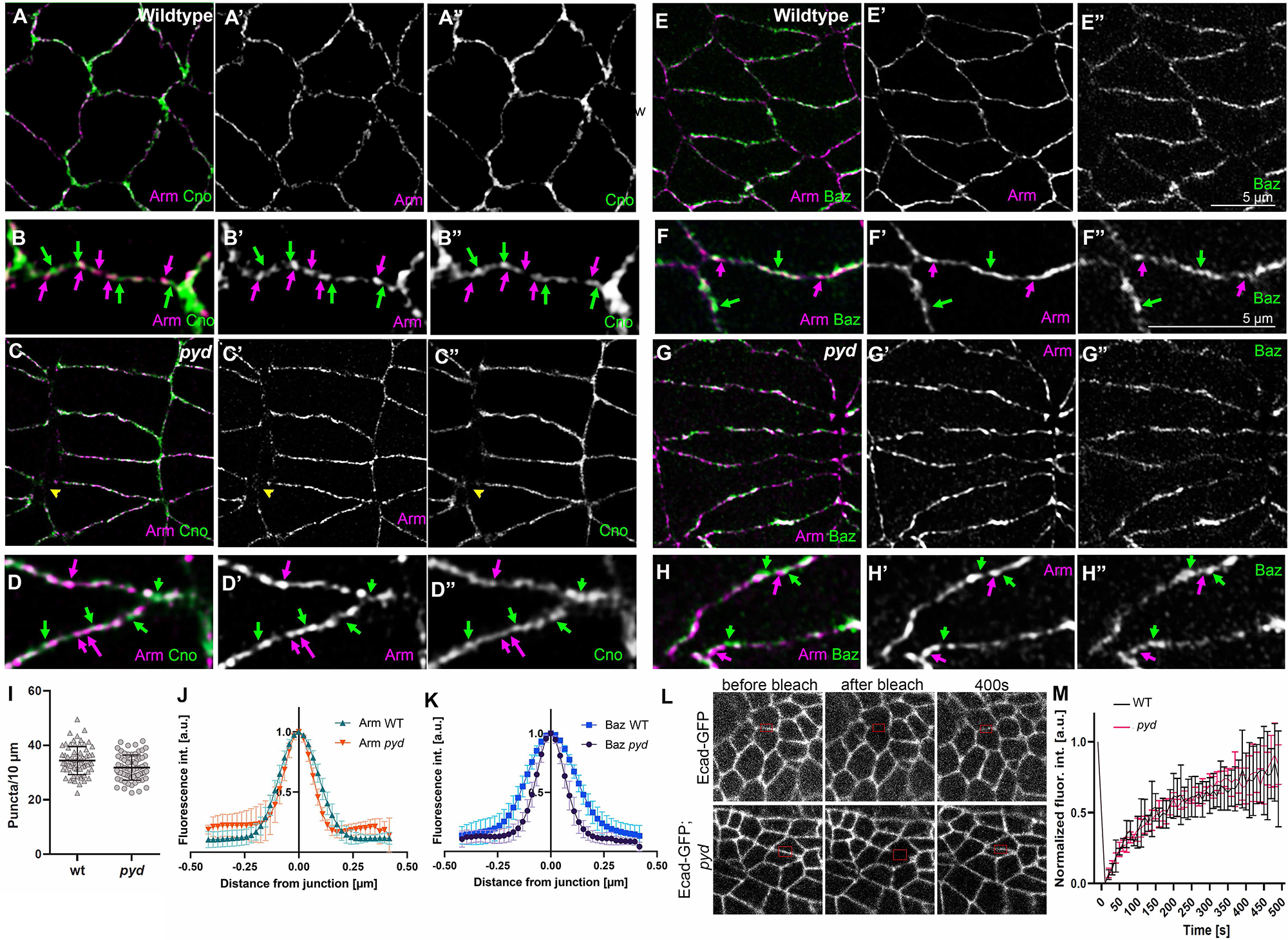
SIM imaging to explore the role of Pyd in AJ molecular architecture. A-H, SIM Imaging, stage 7-8 embryos. A-D. In both wildtype and M/Z *pyd* mutants Arm and Cno continue to differentially localize to puncta along bicellular borders E-H. In both wildtype and M/Z *pyd* mutants Arm and Baz continue to differentially localize to puncta along bicellular borders. I. Arm puncta number per 10µm of bicellular border does not substantially change in M/Z *pyd* mutants relative to wildtype. J. Arm puncta distribution perpendicular to the membrane is not altered in M/Z *pyd* mutants. K. Baz puncta distribution perpendicular to the membrane appears slightly broadened in M/Z *pyd* mutants. M. FRAP of junctional Ecad-GFP in wildtype and M/Z *pyd* mutants. N. Quantification reveals no change in FRAP mobility. Error bars represent SD.

## Discussion

Shaping epithelial tissues and organs during embryonic development requires coordinated cell shape change and cell migration, powered by force generated by the actomyosin cytoskeleton exerted on cell-cell and cell-matrix junctions. As a field, we seek to define the underlying molecular mechanisms. The past two decades saw a growing recognition of the complexity of AJ:cytoskeletal connections that allow cell shape change without disrupting tissue integrity (Yap *et al*., 2018). In this evolving view, simple linear connections mediated by the cadherin-catenin complex were replaced by models involving a robust and multivalent network of proteins. We need to fully define the functions of all the proteins in this network, and to determine the supermolecular structure they assemble at the interface between the cadherin-catenin complex and the cytoskeleton. Here we used *Drosophila* embryonic development as a model, to unravel the function of the scaffold protein Pyd, homolog of human ZO-1, in the robustness of AJ-cytoskeletal connections. In parallel, we used high resolution microscopy to begin to reveal the layered architecture of the AJ in vivo.

### Pyd occupies a new position in the spectrum of proteins mediating AJ:cytoskeletal connections and its phenotypes reveal new complexity in maintaining robustness

In a simple linear model of AJ:cytoskeletal connections, each protein would be equally essential for function. The initial analysis of Ecad, Arm, and alpha-catenin fit this simple model—loss of each totally disrupted cell adhesion, and thus completely halted morphogenesis (Cox *et al*., 1996; Müller and Wieschaus, 1996; Tepass *et al*., 1996; Sarpal *et al*., 2012). Loss of the polarity regulator Baz/Par3 had similar effects (Müller and Wieschaus, 1996; Harris and Peifer, 2004). However, as the field began to assess other players, this simple picture no longer held true. In embryos lacking Cno, for example, cell adhesion is not drastically compromised. However, virtually every morphogenetic movement was disrupted, from mesoderm apical constriction to cell intercalation during germband elongation to collective cell migration during dorsal closure and head involution (Boettner *et al*., 2003; Sawyer *et al*., 2009; Sawyer *et al*., 2011). Further, Cno had mechanosensing properties, responding to elevated tension on AJs, and strengthening AJ-cytoskeletal connections (Yu and Zallen, 2020; Perez-Vale *et al*., 2021). In Cno’s absence AJs and the cytoskeleton detach. These and other data reveal a mechanoresponsive AJ, where force leads to recruitment of multiple proteins to strengthen connections. Surprisingly, many of these proteins are individually dispensable—for example Sidekick, Ajuba, and Vinculin mutant flies are viable and fertile (Maartens *et al*., 2016; Razzell *et al*., 2018; Finegan *et al*., 2019; Rauskolb *et al*., 2019) However, close examination revealed transient defects in AJ:cytoskeletal connections that emerge when cells change shape and move.

We placed Pyd into this continuum. Existing data suggested a protein whose function did not match previously examined. Zygotic null *pyd* mutants are viable, unlike *cno*. However, many (but not all) M/Z *pyd* mutants die as embryos (Choi *et al*., 2011), unlike embryos lacking Sidekick, Ajuba, or Vinculin. We thus directly compared Pyd function to that of Cno in early embryonic morphogenetic events. This revealed that M/Z *pyd* mutant phenotypes overlap with phenotypes of its interaction partner Cno, but that Pyd plays a less essential role. For example, while many M/Z *pyd* mutants have defects in mesoderm invagination, these are often subtle, with the failure to close restricted to the anterior or posterior ends of the mesoderm. Similarly, while we observe apical gaps in AJs under elevated tension—TCJs and aligned AP borders—these are much less frequent than those seen after complete Cno loss, and given the embryonic viability of almost half the M/Z *pyd* mutants, most gaps must re-seal. Finally, the defects in integrity of the ventral epidermis seen after Cno loss are not seen in M/Z *pyd* mutants. Thus, while both Cno and Pyd contribute to making AJ:cytoskeletal connections robust, their relative importance differs.

One of the most striking phenotypes we observed in M/Z *pyd* mutants was alteration in cell shape. In mutants stacks of cells elongated along the AP axis accumulated during germband elongation. Similar phenotypes were observed in *sdk* mutants (Finegan *et al*., 2019), where, as the invaginating hindgut pulls on the tissue, the germband elongates, thus elongating cells, slowed cell rearrangements prevent T1 resolution. The reduced contractility of AJs along the plane of the junction that we observed in M/Z *pyd* mutants may help explain the slowed/stalled constriction of AP borders—*sdk* mutants also have reduced junctional tension (Letizia *et al*., 2019). Similar stacks of elongated cells are also seen in *cno* mutants (Sawyer *et al*., 2011; Perez-Vale *et al*., 2021), but the underlying mechanisms are likely not identical, as the obvious detachment of myosin from AP AJs seen after Cno loss (Sawyer *et al*., 2011) is not seen in M/Z *pyd* mutants. This suggestion of different mechanisms of action is consistent with our earlier observation that Cno and Pyd act in parallel in stabilizing AJs and epithelial integrity— M/Z *pyd* mutants in which Cno is also knocked down using shRNA have defects in AJ stability and epithelial integrity that occur earlier and are much more substantial than those seen in either mutant alone (Manning *et al*., 2019). One way their functions diverge is that Cno is enriched on AP borders, where myosin is enriched and contractility is the highest, while Pyd is enriched with Baz and the cadherin-catenin complex on DV cell borders. However, both are enriched at TCJs, and their respective enrichments at those sites are mutually dependent. This may involve direct protein:protein interactions, which are known to occur between Cno and Pyd (Takahashi *et al*., 1998) and between their mammalian homologs (Ooshio *et al*., 2010). Alternately it may be indirect—for example the reduction in membrane tension seen after Pyd loss might affect Cno recruitment indirectly, via Cno’s mechanosensitive recruitment (Yu and Zallen, 2020; Perez-Vale *et al*., 2021).

Our new data, and the analyses of many labs that have gone before, continue to reinforce and expand the idea that the AJ is a complex, mechanoresponsive machine with many overlapping feedback loops. One example that emerged here is the role of Pyd in maintaining junctional protein planar polarity. M/Z *pyd* mutants resemble *cno* mutants in that AJ proteins and Baz are reduced at AP borders, thus substantially enhancing their planar polarity. Early work defined a reciprocal relationship between Myosin and Baz—they are enriched on opposite cell borders (Bertet *et al*., 2004; Zallen and Wieschaus, 2004), and each antagonizes the others recruitment (Blankenship *et al*., 2006; Simoes Sde *et al*., 2010). Cno loss leads to myosin detachment from AP borders without altering myosin planar polarity (Sawyer *et al*., 2011). Intriguingly, as myosin expands laterally from the AP border, Baz localization becomes more restricted to the center of the DV border, perhaps due to myosin/Baz antagonism. We thus were surprised to see that while Pyd loss strongly elevated Baz planar polarity, it did not disrupt attachment of myosin to AP borders. Thus, altered planar polarity does not require myosin detachment, perhaps suggesting it is a more direct response to contractility.

### High resolution imaging reveals new insights into the complexity of AJ supermolecular architecture

The multiprotein network linking AJs to the cytoskeleton is characterized by proteins that each have multivalent linkages to one another (Fernandez-Gonzalez and Peifer, 2022), raising questions about whether and how they segregate at the architectural level. Scientists studying the cell-matrix junctions pioneered ideas about how a complex network of proteins might form a layered architecture (Case and Waterman, 2015). Work in cultured mammalian cells has begun to suggest a similar layered architecture for cell-cell junctions, with membrane-proximal cadherin tails segregated from the actin cytoskeleton by an interface containing Vinculin, VASP and Zyxin (Oldenburg *et al*., 2015; Bertocchi *et al*., 2017). Electron microscopy also revealed a layered cytoskeletal architecture at bicellular borders, with an Arp2/3-generated branched actin network adjacent to the Cadherin tails, while linear, myosin-decorated actin arrays are found further into the cytoplasm (Efimova and Svitkina, 2018). We sought to extend this effort to examine AJ architecture in vivo during morphogenesis.

In envisioning AJ architecture, we must also consider the accumulating evidence that AJs have at least some attributes of phase-separated biomolecular condensates (Sun *et al*., 2022). These non-membrane bound multiprotein complexes assemble by multivalent interactions, some of which are mediated by intrinsically disordered regions (Harmon *et al*., 2017). Intrinsically disordered regions are a feature of many AJ scaffolding proteins, including Cno/Afadin and Pyd/ZO-1. ZO-1, Pyd’s vertebrate homolog, can phase separate both in vitro and in vivo (Beutel *et al*., 2019; Schwayer *et al*., 2019). ZO-1 droplets can form in the cytoplasm, and as junctions form in cultured cells and zebrafish embryos, they dock on the membrane and then spread. Phase separation of ZO-1 is needed for tight junction formation in 3D cultured cells, and for mechanosensitivity and epithelial tissue spreading during zebrafish gastrulation (Beutel *et al*., 2019; Schwayer *et al*., 2019). Multiple other AJ proteins are also reported to phase separate, including the Baz homolog Par3 (Liu *et al*., 2020) and the Arm homolog beta-catenin (Zamudio *et al*., 2019).

In *Drosophila* AJs assemble during cellularization, as Baz organizes smaller cadherin-catenin complex and Baz puncta into apical spot AJs, with 3-5 SAJs per ∼5µm bicellular border (McGill *et al*., 2009). Each SAJ is roughly 0.5µm in XY and 2.5µm in the XZ, and contains ∼1500 cadherin-catenin complexes and ∼225 Baz proteins (McGill *et al*., 2009). Our data reveal that during germband extension, AJ architecture evolves. At stage 8 we observed 20-50 Arm puncta per 10µm of BCJ—thus puncta number per unit membrane increases 2-5 fold, while average puncta width in the XY was reduced to 0.24µm.

During cellularization Cno localizes with AJ proteins in SAJs, but is more enriched at TCJs—Cno is 5-fold enriched there relative to bicellular SAJs, and at TCJs Cno extends ∼2µm deeper along the apical-basal plane (Bonello *et al*., 2018). Here we used high resolution imaging to examine how the relationship between Cno and the core AJ complex evolves during germband extension. We initially imagined two possibilities: co-localization of Arm and Cno in puncta, or layered architecture, with Cno surrounding Arm. Instead, we got a surprise—while Arm and Cno overlap in many puncta, we observed clear differential localization of these junctional proteins. Their relative enrichment in individual puncta can vary substantially. This was also true for Arm and Baz. In this case the more apical enrichment of Baz relative to Arm was accompanied by differential enrichment in puncta. Our data also suggest that Cno and Baz may extend somewhat more membrane distal than Arm, consistent with Arm’s localization to the membrane-found Ecad tail, while Cno and Baz likely interact in a more multivalent way with multiple junctional and cytoskeletal proteins.

Perhaps the biggest surprise from our imaging was the localization of Pyd. Phase separating ZO-1 can recruit other junctional proteins in vitro (Beutel *et al*., 2019). However, in vivo the localization of Pyd we observed was quite different from that of the other AJ proteins we examined. Rather than localizing to membrane proximal puncta, Pyd localized to strands that overlapped the junctional puncta but could extend much further into the cytoplasm—we observed this with both SIM and Airyscan imaging. Quantification of the breadth of the signal confirmed that Pyd localization to AJs was much broader than Arm, Cno or Baz. One thing that we will ultimately need to consider when interpreting these data is the position of the antigen being visualized and the overall structure of the protein. Pyd, Cno and Baz all contain long intrinsically disordered regions—if these are full extended they could span a considerable distance. Intriguingly, myosin and actin also extend further from the membrane than the core junctional proteins. Together, these data provide a more complex view of AJs during Drosophila germband extension. In one sense AJs are layered, with zones containing different proteins extending different distances from the plasma membrane. However, things are more complex. First, Pyd, actin and myosin extend across the different zones, with Pyd localizing to discrete strands. Second, even within a “layer”, Arm and Cno are differentially enriched in different puncta. Does this imply partial segregation by affinity, with Cno more avidly associating with Cno than with Arm, and vice versa? Future work will be needed to begin to sort out these issues.

### What do these data suggest about Pyd’s mechanism of action?

With our new insights into Pyd localization and the effects of its loss, what might it imply about Pyd’s mechanism of action? We can rule out some simple models. Phase separating ZO-1 can recruit other junctional proteins in vitro, including Afadin (Beutel *et al*., 2019). However, Pyd localization is strikingly different from that of Cno, Baz or Arm. Further, we did not observe obvious mislocalization of Arm, Cno or Baz in M/Z *pyd* mutants. All continued to differentially localize to membrane-proximal puncta, though Baz distribution may have been slightly broadened. This suggests that Pyd is not responsible for the junctional localization of those proteins, or for junctional substructure of the more membrane proximal “layers”. Consistent with this, Pyd did not alter Ecad mobility as assessed by FRAP. Our data also suggest that, unlike Cno, Pyd is not required for the association of myosin cables with AJs along AP cell borders. Pyd does reinforce junctions under elevated tension, but its role is less essential than that of Cno. However, Pyd loss does reduce junctional contractility.

Given this, we think several speculative models are possible. One possibility is that Cno and Pyd play similar but parallel roles, simply linking core junctional proteins and actin, with Cno’s role more essential and with Cno and Pyd differentially strengthening DV versus AP borders. However, their very distinctive localization patterns suggest more substantial differences in function. The Pyd strands extending membrane distal are very intriguing. Mammalian ZO-1 can dimerize via its PDZ2 domain (Utepbergenov *et al*., 2006; Chen *et al*., 2008) and may also dimerize via its SH3/GUK domains (Umeda *et al*., 2006). ZO-1 is critical for formation of the continuous strands of tight junction proteins at mammalian tight junctions (Umeda *et al*., 2006), a function that requires PDZ2 (Rodgers *et al*., 2013). Perhaps Pyd/ZO-1 forms polymers, which we visualize as the strands. We did not see obvious phase separated puncta in the cytoplasm via immunostaining, and the strands we observed are distinct from the more spherical puncta previously seen (e.g. Beutel *et al*., 2019), but these may be present earlier in AJ assembly. It will be of interest to GFP-tag Pyd at its endogenous locus and examine its localization as AJs form and evolve.

We hypothesize that AJs connect to the actomyosin cytoskeleton in different ways at BCJs and TCJs. In this model, the connections of the cytoskeleton to cadherin-catenin clusters at BCJs may not be under the same degree of “molecular force” as those at TCJs. Consistent with this, antibodies detecting the open state of alpha-catenin are enriched at TCJs under tension in mammalian cells (e.g. Choi *et al*., 2016)), and proteins that can sense tension like Cno are enriched at TCJs (Yu and Zallen, 2020; Perez-Vale *et al*., 2021). We imagine that in the germband extending embryo, AJ/cytoskeletal connections are under somewhat elevated tension at AP borders, where contracting myosin cables connect side on to AJs, but that these connections are under even more tension at TCJs, where we envision cables anchor end on to AJs. In M/Z *cno* mutants, myosin detaches and apical junctional gaps arise at both sites (Sawyer *et al*., 2011), suggesting reduced fidelity of both sorts of AJ:cytoskeletal connections. One intriguing aspect of the myosin detachment we previously observed in *cno* mutants is that spot connections to AJs remain, pulled out in strands (Sawyer *et al*., 2011). Perhaps this reflects a role of Cno in lateral cross-linking of cadherin-catenin complexes, allowing them to jointly resist force perpendicular to the membrane. The lack of myosin detachment at AP borders in *pyd* mutants may suggest Pyd does not share this function. Another puzzling issue is why apical AJ gaps appear at aligned AP borders in both mutant genotypes, if force is primarily directed parallel to the membrane in these locations.

We also must define the mechanism by which Pyd maintains tension at AJs. The similar reduction of junctional tension in *sdk* mutants may provide a clue (Letizia *et al*., 2019). Sdk is strongly enriched at TCJs (Finegan *et al*., 2019; Letizia *et al*., 2019), suggesting it acts there. Perhaps both Sdk and Pyd ensure the stability of the hypothesized “end-on” connections of actomyosin cables at TCJs. This role is similar to the role we have suggested Cno plays during dorsal closure, when the leading edge actin cable is anchored cell-cell at specialized leading edge AJs (Manning *et al*., 2019). The localization of the barbed end actin polymerase Ena to TCJs at both stages (Gates *et al*., 2007; Manning *et al*., 2019), and strong genetic interactions between *cno, pyd,* and *ena* (Choi *et al*., 2011) may suggest a special role for anchoring and maintaining actin barbed ends at TCJs. One important experiment remaining is to test whether Cno loss also reduces junctional tension. This is just one of many new questions raised by this work, as the field continues to unravel the complex, multivalent interactions that ensure robust AJ:cytoskeletal connections as cells change shape and move.

## Acknowledgements

We are grateful to Peifer lab members, especially Maik Bischof, for helpful advice and discussions, to the Bloomington Drosophila Stock Centers for fly stocks, and the Developmental Studies Hybridoma Bank for antibodies. We thank Tony Perdue of the Biology Imaging Center for imaging advice and support. This work was supported by NIH R35 GM118096 to M.P. A.S. was supported by DFG (Deutsche Forschungsgemeinschaft) SCHM 3628/1-1. Work in the Grosshans lab is supported by DFG GR1945/15-1, DFG INST1525/16-1 FUGG, and Volkswagenstiftung (A129197, ZN2632) and work in the Wolf lab by DFG FOR1756, GR1945/8-2, GR1945/10-1/2, WO1489/1-2, WO1489/5-1, SFB 1286/C2, the Volkswagenstiftung (A129197, ZN2632) and the Cluster of Excellence Multiscale Bioimaging (MBExC) The authors declare no competing financial interests.

## Author contributions

Anja Schmidt and Mark Peifer conceived the project. Tara Finegan analyzed cell shapes quantitatively and worked with Alex Fletcher to do the modeling. Deqing Kong carried out and analyzed the laser cutting. Matthias Häring carried out the Segmentation and T1 analysis, with help from Anja Schmidt. Zuhayr Alam assisted with genetic analysis. All other experiments were carried out by Anja Schmidt. Jörg Grosshans and Fred Wolf provided funding support and lab space and expertise. Anja Schmidt, Tara Finegan, and Mark Peifer drafted the manuscript, with input from the other authors.

## Material and Methods

### Fly stocks and handling

**Table 1:**
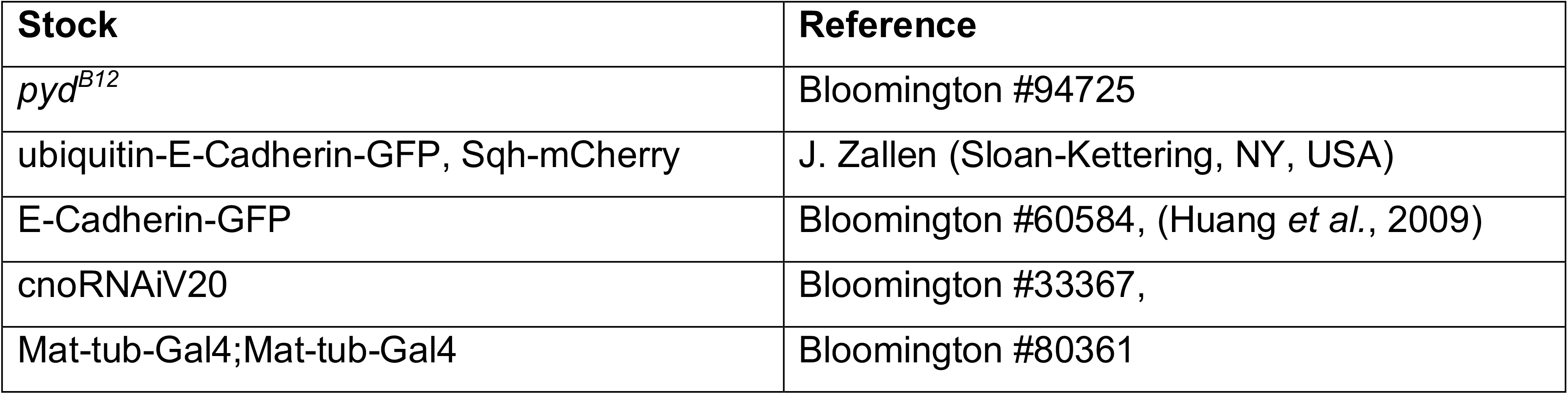
Fly stocks

All crosses and embryo collections were done at 25 °C if not mentioned otherwise. *cno RNAi* embryo collections were kept at 29 °C. *y w* was used as our wild type control. M/Z *pyd* mutant embryos were collected by crossing homozygous mutant virgin females with heterozygous males, as homozygous males are basically sterile. To distinguish M/Z mutant embryos from zygotically rescued ones, the embryos were stained with an antibody directed against Pyd. For live imaging, stocks that were balanced over a TM3 with a GFP-reporter were used.

### Fixation and Immunohistochemistry

Embryos were collected in plastic cups on apple juice agar plates at 25 °C. The embryos were either fixed by formaldehyde treatment or by heat fixation, depending on the staining. For heat fixation, the chorion was removed by treatment with 50 % sodium hypochlorite for 5 min following 3 washing steps with wash buffer (68 mM NaCl, 0.03% Triton). The embryos were then transferred into preboiled salt buffer (68 mM NaCl, 0.03% Triton, 8 mM EGTA) and cooled down after 10 sec with prechilled salt buffer. After the solution cooled down, the embryos were transferred into a 50% heptane, 50% MeOH/EGTA (95% MeOH, 5% EGTA) solution and the vitelline membrane was popped by vigorous shaking, followed by three washing steps in MeOH/EGTA. The embryos were stored at -20 °C in MeOH/EGTA. For formaldehyde fixation, the embryos were collected as above, and after removing the chorion, the embryos were fixed in a 1:1 mixture of heptane and formaldehyde (4% in PBS, 8 mM EGTA) for 20 minutes followed by removal of the vitelline membrane as above. For phalloidin staining and Myosin imaging, fixation was done in 1:1 heptane: 8% formaldehyde in PBS, 8mM EGTA and the vitelline membrane was removed manually.

**Table 2:** Antibodies and probes used.

| Antibodies/probes | Species | Dilution | Source |
| --- | --- | --- | --- |
| Anti-Arm (N2 7A1) | Mouse 2a | 1:100 | Developmental Studies Hybridoma Bank (DSHB) |
| Anti-Baz | Rabbit | 1:1000 | J. Zallen (Memorial Sloan-Kettering Cancer Center) |
| Anti-Cno | Rabbit | 1:1000 | (Sawyer *et al.*, 2009) |
| Anti-Pyd (PYD2) | Mouse 2b | 1:1000 | DSHB |
| Secondary antibodies |  |  |  |
| Alexa Fluor 488, 568, and 647 |  | 1:500 | Life Technologies |
| Other probes |  |  |  |
| Phalloidin-Alexa 488 |  | 1:1000 | Life Technologies |

### Image acquisition

Imaging of fixed samples was performed on the following Zeiss (Oberkochen, Germany) laser scanning confocal microscopes: LSM-5 Pascal with a 40x EC Plan-Neofluar objective (oil, 1.3 NA) or LSM 880 with a 40x Plan-Apochromat objective (oil, 1.3 NA) and 63x Plan-Apochromat objective (oil, 1.4 NA). All fixed and stained embryos were mounted in Aquapolymount or Prolong Diamond mounting media on No. 1.5 cover slides. Airyscan Imaging was done with the Zeiss LSM 880 and Airyscan module with the aforementioned 63x Plan-Aprochromat objective. The pixel size was set to half the size of the theoretical optical resolution resulting in 42.6 nm/pixel and the step size in Z was set to half of the axial resolution resulting in 200 nm. Airyscan processing was done with the automatic settings in ZEN black (Zeiss). SIM imaging was performed on a Nikon N-SIM using a 100x SR APO TIRF objective (oil, 1.49 NA) with a lateral pixel size of 31.8 nm and a step size in Z of 100 nm. Image reconstruction was done with NIS Elements AR Version 4.51 (Nikon).

### Laser ablation and quantification

Laser ablation was conducted as previously described (Lv *et al*., 2022). Briefly, stage 7 embryos from wild type or *pyd* mutant flies expressing Ecad-GFP were collected for junction ablation. Dechorionated embryos were aligned on an agar block and transferred to a cover slide with heptane glue and dried in a desiccation chamber for 2 min, and then covered with halocarbon oil. The UV laser (DPSL355/14, 355 nm, 70 µJ/pulse, Rapp Optoelectronic) was introduced from the epi-port of the confocal microscope (Zeiss LSM 980). Junction ablation was performed with 5% laser power and 50 ms exposure time during the recording mode (100x oil, NA 1.4). The displacement of vertex (L(t)) of ablated junctions was measured manually in Fiji/Image J. The displacement was fitting as a Kelvin-Voigt fiber model (Fernandez-Gonzalez *et al*., 2009) to the following equations on Prism8 (Liang *et al*., 2016)

Extraction of initial recoil values:

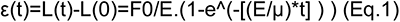

Where, F0 is the tensile force present at the junction before ablation, E is the elasticity of the junction, μ is the viscosity coefficient related to the viscous drag of the cell cytoplasm. As fitting parameters for the above equation initial recoil=dε(0)/dt=F_0/μ (Eq.2) and K values=E/μ (Eq.3) were introduced.

### Live imaging

For live imaging, the embryos were collected from agarose plates and dechorionated in 50% bleach for 90 s and then aligned on a piece of agarose gel and glued to a No. 1.5 cover slide with heptane glue. The embryos were covered with halocarbon oil and either covered by a dish spanned with an oxygen permeable foil or the cover slip was glued to a metal holder with double sided tape.

Live imaging for the analysis of T1 transitions was performed on a Zeiss LSM 980 with Airyscan 2 in multiplex CO-8Y mode with a 63x Plan-Apochromat Oil DIC M27 objective (1.4 NA). The frame size was set to 1564 x 1364 pixel with a lateral pixel size of 85 nm, a step size in z of 0.5 micron, and a frame time of 105.06 ms. We imaged z-stacks in an interval of 15 second, using the multiplex CO-8Y mode with the Airyscan 2 detector.

FRAP was done on a ZEISS LSM 880 and a 63x oil objective as described above. Z-stacks with a step size of 0.5 micron were recorded in an interval of 10 seconds. The frame size was set to 196x196 pixels with a lateral pixel size of 90 nm, an axial step size of 0.5 micron, and a frame time of 121.20 ms. For bleaching, the 488 nm laser was set to 100% laser power with 70 iterations.

### Image analysis and quantification

FRAP of Ecad was done in embryos expressing Ecad-GFP in wild type or *pyd* mutant backgrounds. The analysis was done in Fiji/ImageJ by measuring the mean fluorescence intensity of the bleached region. The axial and lateral movement of the bleached region was corrected manually. To correct for bleaching during imaging, the fluorescence intensity of a junction outside of the bleached region was also measured. Six wildtype junctions in four embryos and seven *pyd* junctions in three embryos were quantified, normalized to the junction outside the bleached region and then normalized to Min and Max to adjust for differences in bleaching before they were averaged. The error bars represent the standard deviation.

For the quantification of the width of Baz, Arm, Cno, and Pyd signal across the junctions, the mean fluorescence intensity of a 10-pixel thick line that was drawn across the junctions was measured in Fiji ImageJ. For all proteins and genotypes a total of nine junctions in three embryos were measured. The individual measurements were then normalized to their maximum which also represented the center on the y axis in the plots. All measurements were averaged, and the error bars represent the standard deviation.

For measurement of cell eccentricity, the ventrolateral regions of fixed embryos stained with an AJ marker were imaged by confocal microscopy and images were processed manually using FIJI/Image J (Schindelin *et al*., 2012) to generate images suitable for automated segmentation. These images were aligned such that the AP axes were oriented left to right. Images were then processed by a custom MATLAB script (https://github.com/tmfinegan/Shape-analysis) to automatically segment cells, and the shape characteristics were extracted using the “regionprops” functionality. Eccentricity is a measure of the ratio of the distance between the foci of an ellipse of best fit and its major axis length. The value is between 0 and 1. An ellipse with eccentricity is 0 is a circle, while an ellipse with eccentricity is 1 is a line segment.

Planar polarity of Baz, Cno and Arm were quantified by line measurements of the mean fluorescence intensity of 20 DV and 20 AP borders in five embryos per genotype was measured. For that, a maximum intensity projection of 2-4 slices that covered the apical zonula adherens was made in Fiji/ImageJ. The line width for the measurements was set to 3 pixels. To correct for background fluorescence or cytoplasmic signal, the mean fluorescence inside 20 cells were measured and subtracted from the border signal. The DV/AP ratios of all embryos were blotted in the graph as well as the average and the error bars represent the standard deviation.

The enrichment of Cno and Pyd at TCJs was quantified in three embryos in in total 30 TCJs per genotype by measuring the mean fluorescence intensity at tricellular junctions and 3 of their surrounding borders. The thickness of the line was set to 3 pixels. Each intensity value of TCJ was then divided by the average of the three neighboring BCJ values. All values were plotted in the graphs and the error bars represent the standard deviations.

For the analysis of apical gaps, they were counted in representative regions of interest (ROIs) of 14 embryos per genotype in stages 7 and 8. Each ROI measured 133 x 133 μm. To qualify as gap, the distance between borders at tricellular- or multicellular junctions had to exceed 1 μm, for long gaps that were aligned at AP borders one gap was counted per 4 cells which corresponds roughly to the number of cells that would make up a rosette. All data points are plotted in the graph with mean and SD.

To count the number of Arm puncta along borders, the spot function in Imaris was used. For automatic detection, the estimated diameter was set to 0.1 μm and the PSF-elongation along the z-axis was set to 0.3 μm, background subtraction was done automatically, and the quality was set above automatic threshold which was afterwards corrected by eye.

Images were processed in Fiji/ImageJ and Adobe Photoshop. 3D images were rendered in Imaris (Oxford Instruments).

### Analysis of T1 transitions

For the segmentation of germband tissue, the 3D data was reduced to 2D by summarizing the fluorescence intensities in ZEN black. We used a custom designed image segmentation pipeline to segment the tissue. It’s core element is the machine learning based skeletonizing of the tissue, using cycle-consistent generative adversial networks (CycleGAN) (Häring *et al*., 2018). The predicted segmentation was then manually corrected, using the Tissue Analyzer plugin in Fiji/ImageJ (Aigouy *et al*., 2016). Afterwards, the data was parsed via TissueMiner (Etournay *et al*., 2016), and analyzed with custom written software in Python.

The detection of T1 transition was achieved by assignment of neighbor exchanges, which were defined by loss of contact of two cells with gaining new cell contacts to cells which were previously neighboring the initial cell pair. To be included in the quantification, we required a successful tracking over 4 frames as well as a successful T1 event was defined by maintenance of the newly established junction for at least 4 frames and the new junction had to obtain a minimal threshold length of 1 micron. The cell quadruplet must be stable and retain its neighbor connections for at least 10 frames. Furthermore, the area of the cells within the quadruplet was not allowed to change more than 20%.

#### T1 orientation

The orientation of a T1 transition is obtained by computing a *nematic director* (Etournay *et al*., 2016) from the midpoints of the participating cells. The orientation takes values in (-pi/2, pi/2), which is the angle with respect to the experimentally determined anterior-posterior axis of the tissue.

#### 4x points

The 4x points are the events where two cells of the T1 quadruplet are losing contact. If the new junction is immediately established, we speak of a direct T1 transition. The 4x point is called meta-stable if it takes time to resolve into establishing a new junction. The last 4x point is the one right before successful junction establishment.

#### Rate of T1 resolution

The T1 resolution rate is determined by fitting an exponential distribution to the cumulative distribution function of T1 attempt duration. The rate of the exponential, which equals the inverse of the mean, is called the rate of T1 resolution. Intuitively, it quantifies how long the resolution of a T1 transition will take on average. Smaller rates correspond to a longer expected attempt duration.

#### Confidence intervals

If not stated otherwise, error regions have been determined by calculating 95% bias-corrected and accelerated bootstrap confidence intervals with 5000 samples.

#### Statistical tests

If not stated otherwise, the *p* values for estimators have been calculated from one- or two-sided t-tests. The *p* values to compare distributions were calculated using Kolmogorov-Smirnov tests.

### Vertex model of axis extension in wild type and *pyd* mutant

We used mathematical modelling to investigate the topological implications of actomyosin contractility during *Drosophila* axis extension in a *pyd* mutant, modifying a vertex model of wild type behavior (Tetley *et al*., 2016). Vertex models describe epithelial mechanics by considering the polygonal tessellation that cells’ adherens junctions form in two dimensions (Fletcher *et al*., 2014). In such models, the movement of junctional vertices and the intercalation of cells are governed by the strength of cell-cell adhesion, the contractility of the actomyosin cortex and cell elasticity. We describe the epithelial sheet by a set of connected vertices in two dimensions. Assuming that the motion of these vertices is overdamped, the position *r_i_*(*t*) of vertex *i* evolves according to the first-order equation of motion

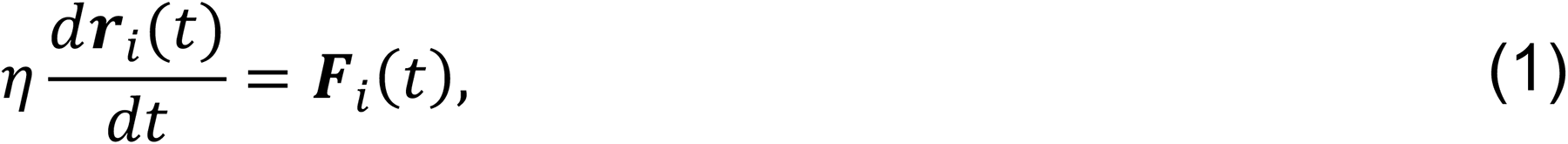

where ***F**_i_*(*t*) denotes the total force acting on vertex *i* at time *t* and η denotes the common drag coefficient. We specify the forces acting on vertices through a ‘free energy’ function *U*, for which

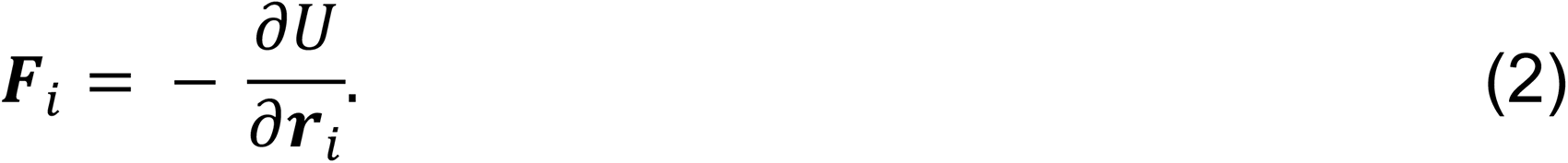

Our choice of *U* is given by (Farhadi *et al.al.*, 2007).

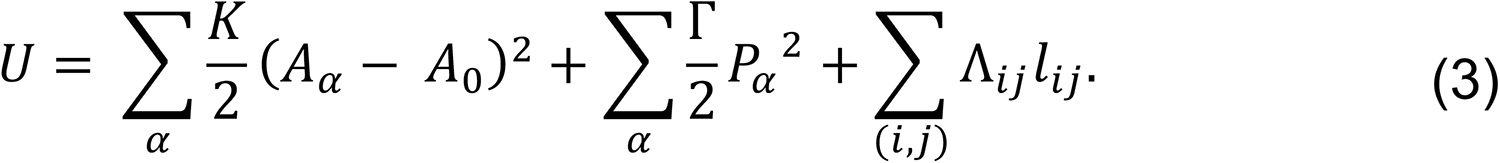

The first term in this free energy function describes an area elasticity with common elastic coefficient *K*, for which *A*_α_ is the area of cell α and *A*_6_ is a common ‘target’ area, and the sum runs over all cells at time *t*. The second term describes the contractility of the cell perimeter *P_α_* by a common coefficient Γ, with the sum again running over all cells at time *t*. The third term represents ‘line tensions’ at cell-cell interfaces, where *l_ij_* denotes the length of the interface shared by vertices *i* and *j*, the line-tension coefficient Λ_*ij*_ may take different values for different edges, and the sum runs over the set of interfaces at time *t*. Line tensions can be increased by reducing cell-cell adhesion and/or increasing junctional myosin activity. The precise functional form of this line tension term varies across our simulations.

In addition to these equations of motion for cell vertices, we must ensure that cells are always non-intersecting and allow cells to form and break bonds. This is achieved through an elementary operation called edge rearrangement (a ‘T1 swap’), which corresponds biologically to cell intercalation. Mathematically, such arrangements are necessary in the vertex model due to the finite forces acting on a cell’s vertices arbitrarily far from equilibrium. We implement a T1 swap whenever two vertices *i* and *j* are located less than a minimum threshold distance *d_dim_* apart (taken to be much smaller than a typical cell diameter). In this case, the two vertices are moved orthogonally to a distance *pd_dim_* apart and the local topology of the cell sheet is modified such that they no longer share an edge.

We explicitly model the presence of an extrinsic pull in the posterior (horizontal; positive *x*) direction, representing the action of the posterior midgut. This extrinsic pull is modelled as follows. At each time step, after moving each vertex a small amount according to its equation of motion and executing any neighbor exchanges (see below), we manually displace each vertex horizontally by an amount that scales linearly with the vertex’s distance from the anterior edge (minimum *x*) of the tissue. This scaling is chosen so that the extrinsic pull exhibits a constant strain rate across the tissue. In addition, we ‘pin’ the anterior edge of the tissue by preventing the anterior-most vertices to move in the *x* direction, though allowing them to move in dorsoventrally (vertically; *y* direction).

In summary, the configuration of the tissue is updated using the following algorithm. Starting from an initial configuration *r*_i_(0), we update the state of the system until time *T* over discrete time steps Δ*t*. At each time step we: implement any required T1 swaps; compute the forces ***F***_i_ on each vertex from the free energy *U*; solve the equation of motion for each vertex over the time step numerically, using an explicit Euler method; implement an extrinsic pull and boundary conditions, if specified; and finally update the positions of all vertices simultaneously. We implement this model in Chaste, an open source C++ library that allows for the simulation of vertex models (Fletcher *et al*., 2013).

We introduce four distinct ‘stripes’ of cell identities within each parasegment (Tetley *et al*., 2016). In our model, the line-tension coefficient Λ_ij_ takes one of three values, depending on the type of interface: (i) if the interface is shared by two cells of the same identity, or belongs to a single cell, then Λ_ij_ = Λ_int_; (ii) if the interface is shared by two cells whose identities (modulo 4) differ by 1, then Λ_ij_= Λ_bdy_; (iii) if the interface is shared by two cells whose identities (modulo 4) differ by 2 (‘mismatched boundary interface’), then we set Λ_ij_ = Λ_*sub*_. We refer to (i)-(iii) as ‘internal’, ‘boundary’ and ‘mismatched boundary’ interfaces, respectively. We choose Λ_*int*_< Λ_bdy_ < Λ_sub_ to reflect our assumption that boundary interfaces are more contractile than internal interfaces, and mismatched boundary interfaces are even more contractile.

We model the movement, shape change and neighbor exchange of a small tissue that is initially comprised of 20 rows and 14 columns of hexagonal cells. Prior to the start of each scenario, we simulate the evolution of the tissue to mechanical equilibrium under the assumption that the line-tension coefficient takes the same value for every interface, Λ_ij_ = Λ_*int*_. This avoids compounding the later dynamics by ‘artifacts’ associated with starting the tissue away from the equilibrium cell size. The value of Λ_*int*_, and all other parameters used in the scenarios described below, are provided in Table 1. We then modify the line-tension coefficients to take different values for internal, boundary and mismatched boundary interface, and implement an extrinsic pull on the tissue, as described above. Under these conditions we simulate the tissue for some time *T*.

**Table 3:** The parameters and their values used in our vertex model simulations.

| Parameter | Description | Value | Reference(s) |
| --- | --- | --- | --- |
| $\eta$ | Drag coefficient | 1.0 | (Kursawe *et al.*, 2017) |
| $T$ | Simulation end time | 300 | - |
| $\Delta t$ | Time step | 0.001 | (Kursawe *et al.*, 2017) |
| $d_{min}$ | T1 swap threshold | 0.01 | (Kursawe *et al.*, 2017) |
| $p$ | T1 swap distance multiplier | 1.5 | (Kursawe *et al.*, 2017) |
| $K$ | Elastic coefficient | 1 | (Farhadifar *et al.*, 2007) |
| $A_0$ | Cell target area | 1 | (Farhadifar *et al.*, 2007) |
| $\Gamma$ | Contractility coefficient | 0.04 | (Farhadifar *et al.*, 2007) |
| $\Lambda_{int}$ | Internal line-tension coefficient | 0.05 | (Tetley *et al.*, 2016), |
| $\Lambda_{bdy}$ | Boundary line-tension coefficient | $2\Lambda_{int}$ | (Tetley *et al.*, 2016), |
| $\Lambda_{sup}$ | Mismatched boundary line-tension coefficient | $8\Lambda_{int}$ | (Tetley *et al.*, 2016), |

---

**Figure S1:**
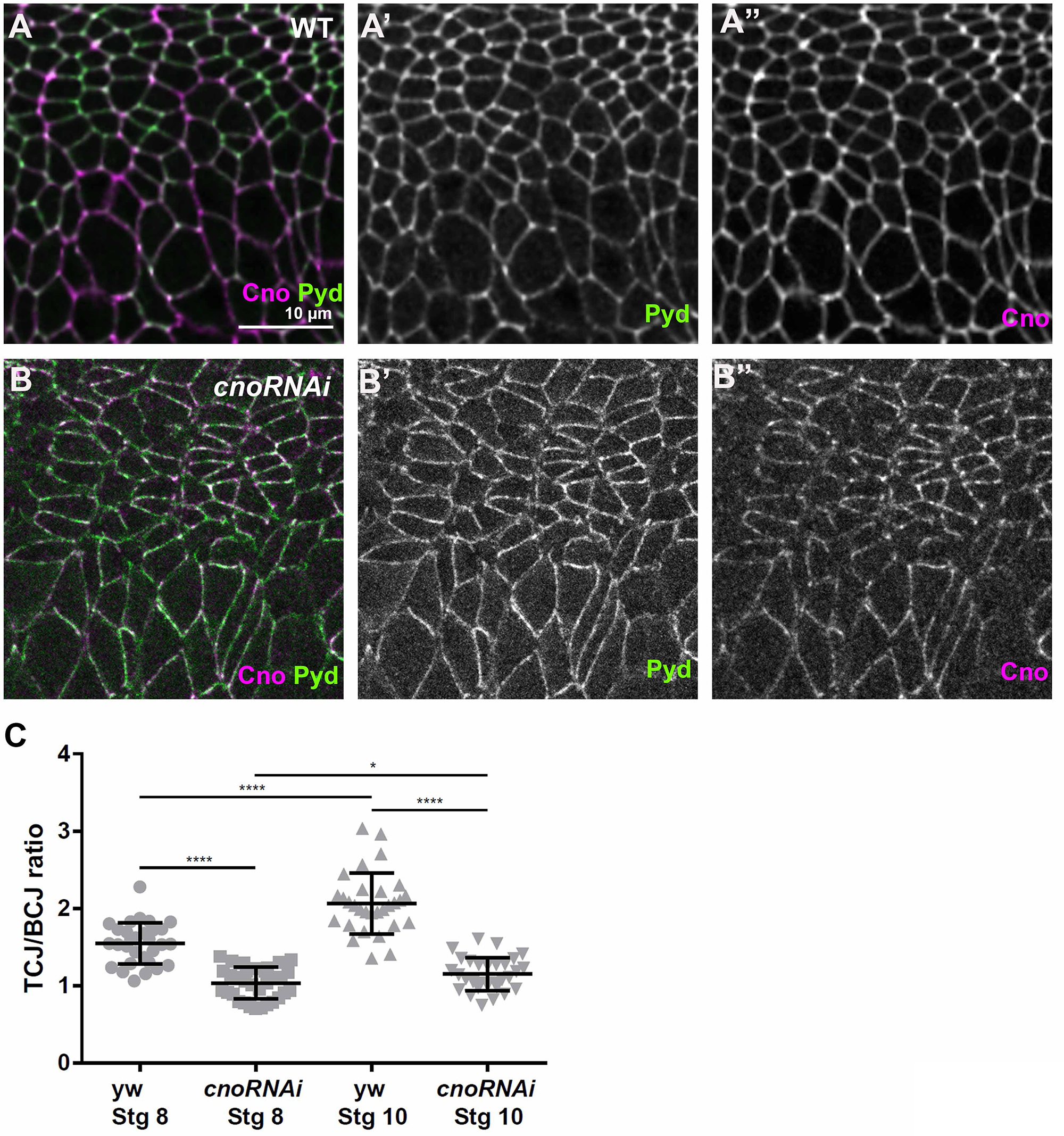
TCJ enrichment of Pyd is further elevated during later embryogenesis. (A-B) Fixed stage 9-10 wildtype (WT) (A) or *cnoRNAi* (B) embryos stained for Pyd (green) and Cno (magenta). (C) Quantification of the ratio of fluorescence intensity at TCJ versus BCJ at stages 8 and stages 9-10 as indicated. Error bars represent standard deviation and p-values are as follows * ≤ 0.05, ** ≤ 0.01, *** ≤ 0.001, **** ≤ 0.0001.

## Notes

### Competing Interest Statement

The authors have declared no competing interest.

